# Safe focused ultrasound-mediated blood-brain barrier opening is driven primarily by transient reorganization of tight junctions

**DOI:** 10.1101/2025.01.28.635258

**Authors:** Rebecca Lynn Noel, Tara Kugelman, Maria Eleni Karakatsani, Sanjid Shahriar, Moshe J. Willner, Claire Sunha Choi, Yusuke Nimi, Robin Ji, Dritan Agalliu, Elisa E. Konofagou

## Abstract

Focused ultrasound (FUS) with microbubbles opens the blood-brain barrier (BBB) to allow targeted drug delivery into the brain. The mechanisms by which endothelial cells (ECs) respond to either low acoustic pressures known to open the BBB transiently, or high acoustic pressures that cause brain damage, remain incompletely characterized. Here, we use a mouse strain where tight junctions between ECs are labelled with eGFP and apply FUS at low (450 kPa) and high (750 kPa) acoustic pressures, after which mice are sacrificed at 1 or 72 hours. We find that the EC response leading to FUS-mediated BBB opening at low pressures is localized primarily in arterioles and capillaries, and characterized by a transient loss and reorganization of tight junctions. BBB opening still occurs at low safe pressures in mice lacking caveolae, suggesting that it is driven primarily by transient dismantlement and reorganization of tight junctions. In contrast, BBB opening at high pressures is associated with obliteration of EC tight junctions that remain unrepaired even after 72 hours, allowing continuous fibrinogen passage and persistent microglial activation. Single-cell RNA-sequencing of arteriole, capillary and venule ECs from FUS mice reveals that the transcriptomic responses of ECs exposed to high pressure are dominated by genes belonging to the stress response and cell junction disassembly at both 1 and 72 hours, while lower pressures induce primarily genes responsible for intracellular repair responses in ECs. Our findings suggest that at low pressures transient reorganization of tight junctions and repair responses mediate safe BBB opening for therapeutic delivery.

**Significance Statement:** Focused ultrasound with microbubbles is used as a noninvasive method to safely open the BBB at low acoustic pressures for therapeutic delivery into the CNS, but the mechanisms mediating this process remain unclear. Kugelman et al., demonstrate that FUS-mediated BBB opening at low pressures occurs primarily in arterioles and capillaries due to transient reorganization of tight junctions. BBB opening still occurs at low safe pressures in mice lacking caveolae, suggesting a transcellular route-independent mechanism. At high unsafe pressures, cell junctions are obliterated and remain unrepaired even after 72 hours, allowing fibrinogen passage and persistent microglial activation. Single-cell RNA-sequencing supports cell biological findings that safe, FUS-mediated BBB opening may be driven by transient reorganization and repair of EC tight junctions.

## Introduction

Targeted delivery of therapeutics to the central nervous system (CNS) to treat neurological and psychiatric diseases remains a challenge due to the presence of a neurovascular barrier, termed the blood-brain barrier (BBB), that protects the CNS from potentially damaging substances, antibodies and immune cells.^1,2^ The BBB is characterized by a low rate of vesicular transport due to a small number of endocytosis vesicles, termed caveolae, compared to the non-CNS vasculature, and a high trans-endothelial electrical resistance mediated by tight junctions (TJ) formed between endothelial cells (ECs). Membrane transporter proteins in brain ECs control the uptake of nutrients and efflux of waste and potential toxins across the BBB. Tight regulation of caveolae-mediated transcytosis^3–5^ and the presence of tight junctions (TJs), forming a charge- and size-selective barrier, limit the transport of molecules across BBB; yet, these cell biological properties of the CNS vasculature pose a hurdle for drug delivery.^1,2^

Over the past two decades, neurosurgical, pharmacological, and physiological strategies have been developed to circumvent the BBB. Focused ultrasound with systemically delivered microbubbles (<10 µm in diameter; FUS) is a technology that increases BBB permeability locally, reversibly and non-invasively to facilitate targeted CNS drug delivery.^6^ The emitted ultrasonic wave, propagating though the intact scalp and skull, causes microbubbles only within the focal zone of the FUS beam to oscillate, leading to cavitation within the vasculature. At low acoustic pressures, the cavitating microbubbles undergo stable volumetric oscillations, termed stable cavitation, whereas at high acoustic pressures the microbubbles may experience inertial cavitation resulting in microstreaming and shock waves.^7–11^ Multiple studies have optimized the frequency,^12^ acoustic pressure,^13–15^ pulse duration,^16,17^ pulse and burst repetition frequency^18^ and microbubble size ^19,20^ used to open the BBB in a safe and effective manner in rodents, non-human primates^21,22^ and humans.^23,24^ Using the FUS microbubble technology, numerous chemotherapeutic agents,^25–27^ antibodies,^28,29^ neurotrophic factors,^30,31^ adeno-associated viruses, ^32,33^ and neural stem-cells^34^ have been delivered across the BBB in several neurological diseases.

Although parameters for safe effective opening of the BBB in rodents, non-human primates, and humans have been worked out, the molecular mechanisms mediating BBB opening and the response of the neurovascular unit (NVU) to FUS-mediated BBB breakdown at either a low acoustic pressure associated with stable cavitation and safe BBB opening, or a high acoustic pressure associated with inertial cavitation that causes tissue damage, remain poorly characterized. Previous transmission or immunoelectron microscopy studies have shown that FUS with a high acoustic pressure (1MPa), 1.5 MHz or 1.63 MHz center frequency transducer, and 100 ms pulse duration increases the number of caveolae and fenestrae within the CNS endothelium leading to formation of trans-endothelial channels to transport therapeutics across the BBB.^35^ Similarly, high acoustic pressures (1 MPa) and shorter pulse durations (10 ms) with the same 1.5 MHz center frequency transducer induce loss of endothelial TJ-associated proteins by immune EM within one hour after FUS administration, suggesting a potential increase in paracellular BBB permeability.^36^ In contrast, at pressures ranging from 200 to 800 kPa using a 1.029 MHz and 1.2 MHz center frequency transducer, two-photon imaging studies have described two distinct rates of diffusion across the BBB: a slow diffusion primarily found in larger caliber vessels at a rate comparable to the transcellular transport, and a quick diffusion in smaller caliber vessels at a rate comparable to paracellular diffusion likely through disrupted tight junctions.^37,38^ Thus, FUS can change both transcellular and paracellular BBB permeability depending on the applied pressure.

Despite these findings, the temporal cell biological changes in BBB properties and the molecular mechanisms driving these changes at either low acoustic pressure that opens transiently the BBB, or high pressure that causes brain damage, remain unclear since the FUS parameters change significantly among studies. In addition, the cellular response of other NVU cells such as microglia or astrocytes after FUS-mediated BBB opening remains poorly characterized. In this study, we examined endothelial TJ morphology in CNS blood vessels of *Tg::eGFP-Claudin5* transgenic mice where TJs are labelled with eGFP^5,39^ at two distinct pressures and time points to correlate temporal changes in FUS-mediated BBB properties with structural TJ alterations. We have chosen two pressures, a 450 kPa pressure associated with stable cavitation and a safe and reversible BBB opening, and a 750 kPa one associated with inertial cavitation and known to cause irreversible damage.^40,41^ We have selected two time-points: a) 1 hour post FUS, when the BBB permeability is increased at both low and high pressures, and b) 72 hours post FUS, when the BBB is resealed at low pressure, but remains permeable at high pressure.^40,41^ We report that FUS-mediated BBB opening is associated with permanent damage of endothelial TJs in both arterioles and capillaries by confocal microscopy within 1 hour only at high pressure; these junctions remain unrepaired even at 72 hours. However, venules are excluded from damage demonstrating vessel selectivity for FUS-mediated BBB opening. This allows fibrinogen passage at sites of BBB leakage leading to persistent microglial activation. In contrast, at low safe pressures, BBB opening still occurs due to transient loss and reorganizations of TJs in arterioles and capillaries of both wild-type and *Cav1*^-/-^ mice, suggesting a TJ-dependent, but caveolar-independent, mechanism for transient BBB impairment. Single-cell RNA sequencing of arteriole, capillary and venule ECs reveals that the transcriptomic responses of cells exposed to high FUS pressure at both 1 and 72 hours are dominated by genes belonging to the stress response and junction protein disassembly, while lower pressures primarily drive genes responsible for intracellular repair responses in ECs. These findings suggest that low pressures can safely and transiently open the BBB to deliver therapeutics to the brain primarily through transient reorganization of tight junctions.

## Results

### FUS with microbubbles increases BBB permeability locality in *Tg::eGFP-Claudin5^+/-^* mice

To examine the relationship between the temporal changes in BBB permeability due to FUS with microbubbles (denoted as FUS throughout the paper) with endothelial TJ morphology by confocal microscopy, we used a stereotactic system to non-invasively target the focused ultrasound beam to the left caudate and putamen brain regions of *Tg::eGFP-Claudin5^+/-^* mice, where TJs are labelled with eGFP.^5,39^ Immediately preceding sonication, mice were injected intravenously with microbubbles and biocytin-TMR, a small (∼890 Da) fluorescent tracer to assess BBB permeability.^5,39,42,43^ FUS-mediated BBB opening was performed at either 450 kPa, a low pressure that can safely open the BBB and deliver various agents into the CNS,^40,44^ or 750 kPa, a high pressure known to cause irreversible CNS damage^40,41^ (**Figure 1A**). Throughout the sonication, passive cavitation detection signals were collected and cumulative stable and inertial cavitation doses were calculated for each pressure (**Figure S1**). Harmonic acoustic signals associated with stable cavitation^7^ were detected at both pressures (**Figure S1A, B)**. At high pressure (750 kPa), ultra-harmonics^45^ and broadband emissions were also detected (**Figure S1B, D, F**), indicative of inertial cavitation.^7^ The stable cavitation dose for harmonics was not significantly different between 450 and 750 kPa (**Figure 1B**); however, the doses for ultra-harmonic and inertial cavitation were significantly different between two pressures (**Figure 1B**).

**Figure 1.**
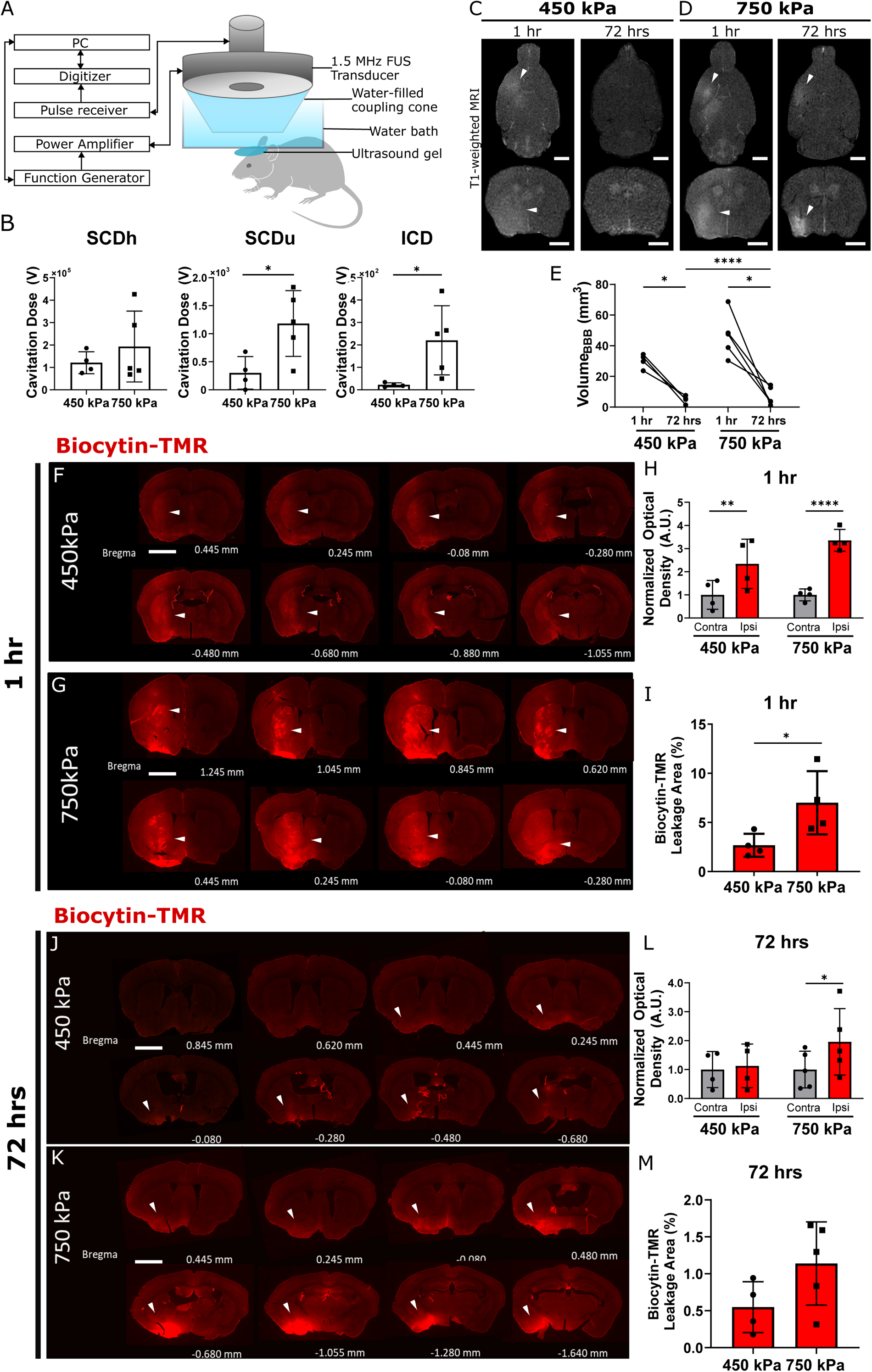
FUS with microbubble induces dynamic transient BBB opening only at safe pressures. (**A**) Schematic diagram showing a 1.5 MHz focused ultrasound transducer targeting the caudate/putamen region of the brain through the scalp and skull. (**B**) Quantification of the stable cavitation dose (SCD) for harmonics (SCDh) and ultra-harmonics (SCDu), along with the inertial cavitation dose (ICD) for 450 kPa (N=4 mice) and 750 kPa (N=5 mice). Signals were collected from a passive cavitation detector (PCD) during a 120 second interval of sonication. (**C-D**) Axial and coronal evaluation of BBB opening at 45 min and 71 hours and 45 min post FUS with contrast-enhanced T1-weighted MRI in the caudate / putamen region for 450 kPa (**C**) and 750 kPa (**D**) pressures. The BBB becomes permeable at both pressures after 1 hour; however, the BBB is permeable only at 750 kPa after 72 hours. (**E**) Quantification of the BBB leakage volume at 45 min following gadolinium injection administered immediately after FUS and at 71 hours post FUS (N=4 mice/group; *p<0.05 and ****p<0.0001; two-way ANOVA with Sidak’s multiple comparison test). (**F, G**) *Tg::eGFP-Claudin5*^+/-^ mice were injected with biocytin-TMR (870 Da) tracer after 1.5 Mhz frequency FUS to determine BBB permeability. Brain sections from *Tg::eGFP-Claudin5*^+/-^ mice showing biocytin-TMR tracer leakage from the vasculature on the ipsilateral caudate/putamen region 1 hour following FUS-mediated BBB opening at both 450 kPa and 750 kPa. (**H**) Quantification of biocytin-TMR fluorescence intensity normalized to optical density between ipsilateral and contralateral matched regions of interest (ROIs) at 450 kPa and 750 kPa pressures 1 hour after FUS (N=4 mice/group; **p<0.01 and ****p<0.0001; two-way ANOVA with Sidak’s multiple comparison test; a.u., arbitrary unit). (**I**) Quantification of biocytin-TMR leakage area at 450 kPa and 750 kPa one hour after FUS (N=4 mice/ group; *p<0.05; Student’s t-test). (**J-K**) Brain sections from *Tg::eGFP-Claudin5*^+/-^ mice showing biocytin-TMR tracer leakage in the caudate/putamen region 72 hours following FUS-mediated BBB opening at 450 kPa (N=4) and 750 kPa (N=5). There is no biocytin-TMR in the brains of mice that received FUS at low pressure (450 kPa). However, choroid plexus that has a leaky vasculature is intensity labelled with biocytin-TMR. (**L**) Quantification of biocytin-TMR fluorescence intensity normalized to optical density between ipsilateral and contralateral matched ROIs at 450 kPa (N=4) and 750 kPa (N=5) 72 hours after FUS-mediated BBB opening (*p<0.05; two-way ANOVA with Sidak’s multiple comparison test; a.u., arbitrary unit). (**M**) Quantification of biocytin-TMR leakage area at 450 kPa (N=4) and 750 kPa (N=5) 72 hours after FUS. Student’s t-test was used for statistical comparisons (p=0.1106). All data are presented as mean± standard deviation. Arrowheads for Figure 1F, G, J and K denote the area of biocytin extravasation. See also **Figure S1**.

Acute and long-term BBB leakage was assessed with contrast enhanced T1-weighted magnetic resonance imaging (MRI). Gd-DPTA-BMA (∼540 Da) was injected intraperitoneally either immediately following FUS or 71 hours post-FUS and scans were acquired after 45 min to allow for tracer diffusion. At both time points and pressures, diffusion of Gd-DPTA-BMA across the BBB was observed within the sonicated hemisphere (**Figure 1C-D**). Contrast enhancement was significantly greater at 1 compared to 72 hours for both pressures, and at 750 kPa compared to 450 kPa by 72 hours (**Figure 1E**). Signal enhancement was significantly greater in mice exposed to high compared to low pressure at 72 hours, suggesting restored BBB function at low pressure and sustained BBB damage at high pressure.

To further characterize the functional changes in BBB permeability, we examined the diffusion of biocytin-TMR (890 Da) across blood vessels into the brain parenchyma in tissue sections one hour following FUS sonication with 450 or 750 kPa (**Figure 1F, G**). The normalized optical density (NOD) of biocytin-TMR was 2- and 3.5-fold higher in the ipsilateral compared to the contralateral hemisphere 1 hour after FUS at low and high pressure, respectively, (**Figure 1H**). The area of biocytin-TMR leakage was 4-fold higher in mice sonicated at 750 kPa compared to 450 kPa within 1 hour after FUS (**Figure 1I**). The rate of BBB closure highly depends upon the applied FUS peak rarefactional pressure.^40^ We assessed whether BBB permeability was restored by 72 hours after FUS application by measuring biocytin-TMR diffusion across the vessels at both pressures. The NOD was not significantly different between ipsilateral and contralateral hemispheres at 450 kPa, suggesting effective BBB closure (**Figure 1J, L**); however, it was 2-fold greater on the ipsilateral hemisphere compared to the contralateral at 750 kPa, indicating sustained BBB disruption without closure at high pressure (**Figure 1 K, L**). The brain area permeable to biocytin-TMR at 72 hours after FUS was higher, albeit non-significant, in mice exposed to 750 kPa compared to 450 kPa pressures (**Figure 1M**), in contrast to the Gd-DPTA-BMA permeability at 72 hours (**Figure 1E**). Overall, these findings demonstrate a significantly higher BBB permeability at 72 hours in high compared to low FUS pressure. Moreover, the BBB opens transiently only at low (safe) acoustic pressures but remains permeable at higher (damaging) pressures over time.

### Loss of tight junctions in both arterioles and capillaries contributes to persistent FUS-mediated BBB opening at high acoustic pressures

To correlate changes in FUS-mediated BBB permeability at both pressures with structural alterations in TJs, we analyzed the number and type of leaky vessels as well as structural TJ abnormalities via immunofluorescence and confocal imaging in the ipsilateral and contralateral caudate putamen regions at 1 and 72 hours after FUS. We assessed the spatial localization of biocytin-TMR relative to a blood vessel marker, glucose transporter 1 (Glut-1/Slc2a1). The majority (∼90%) of leaky vessels were small (<10 µm) than large (>10 µm) diameter vessels for both pressures at 1 hour after FUS (**Figure 2A**). There was a 3-fold increase in the number of leaky Glut1^+^ small diameter vessels (<10 µm) and a 4-fold increase in the number of leaky large diameter vessels (>10 µm) at high compared to low pressures after 1 hour (**Figure 2A**). There was no significant difference between the percentage of small versus large diameter leaky vessels at 450 kPa, albeit larger diameter vessels seemed more affected (**Figure 2B**). In contrast, a greater percentage of large, compared to small, diameter vessels were significantly leaky at 750 kPa (**Figure 2B**). The percentage of leaky large vessels was greater at 750 compared to 450 kPa (**Figure 2B**). These data indicate that higher pressures increase the probability of opening larger diameter vessels. The larger leaky vessels were exclusively arterioles based on endothelial TJ morphology^5,39^ and expression of α-smooth muscle actin (α-SMA; data not shown). We could not detect tracer leakage from venules or larger veins (data not shown). By 72 hours, the average number of leaky vessels at both pressures was lower than 1 hour post-FUS, and the number of leaky small vessels was higher than large ones (**Figure 2C**). However, the vessel diameter was a poor predictor of the area and intensity of tracer leakage at either pressure 1 hour after FUS (**Figure S2A-B, E-F**). To assess whether small or large caliber vessels are more likely to remain permeable 72 hours after FUS, we quantified the percentage of each leaky vessel type in the caudate putamen region. There was no significant difference between the percentage of leaky small vs. large diameter vessels in each pressure; however, the percentages of leaky small and large diameter vessels were greater at high compared to low pressure (**Figure 2D**). The percentage of leaky small or large vessels remained high for 750 kPa at 72 hours (**Figure 2D**), consistent with biocytin-TMR leakage across the BBB (**Figure 1J-M**). These data suggest that low safe pressure does not preferentially disrupt smaller compared to larger diameter vessels, and this disruption is largely resolved by 72 hours. In contrast, large vessels are preferentially disrupted at high pressures, and both small and large diameter vessels remain leaky 72 hours after FUS.

**Figure 2.**
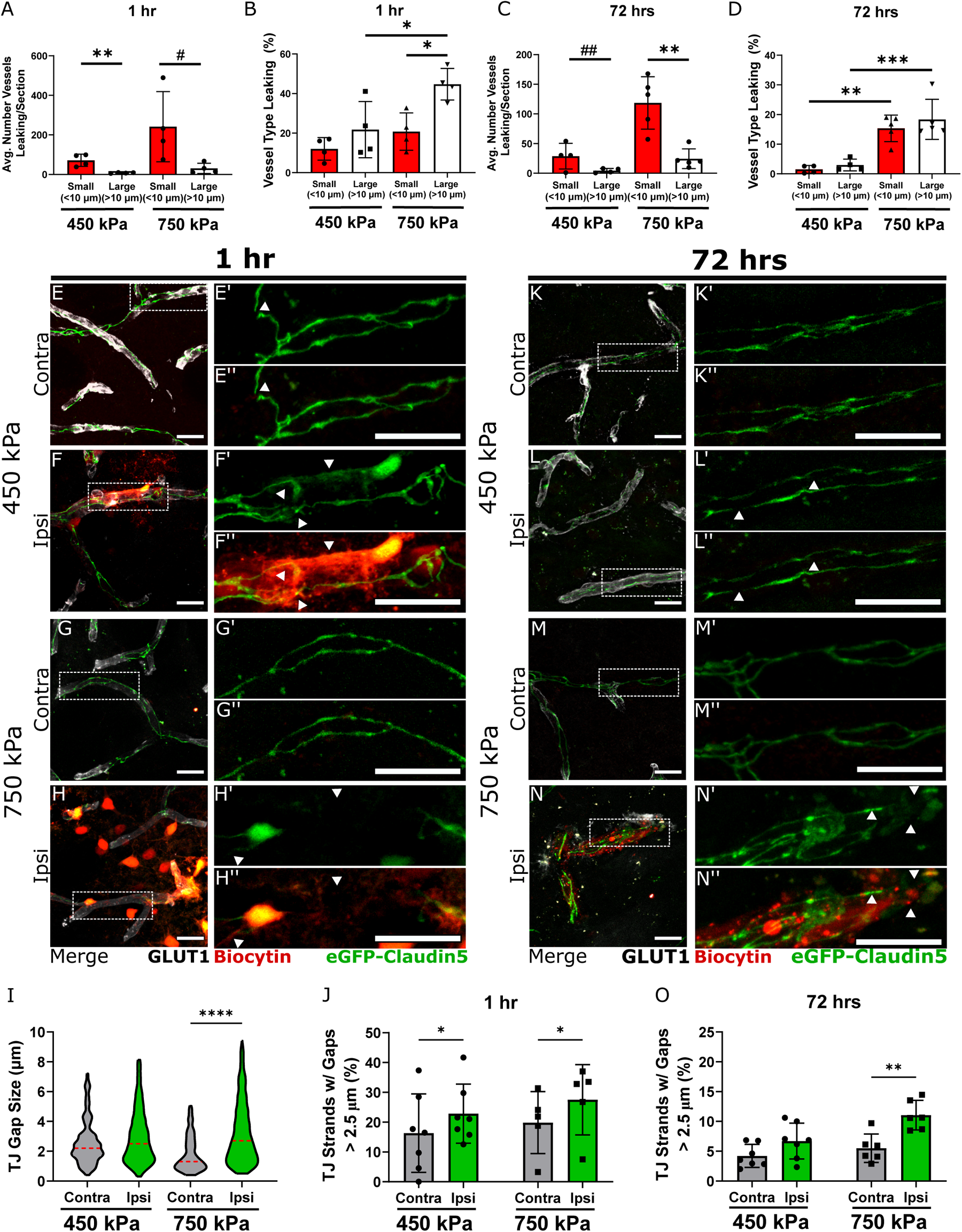
Persistent structural abnormalities in capillary tight junctions visualized by confocal microscopy correlate with continuous BBB opening at unsafe pressures. (**A**) Quantification of the average number of leaky small (<10 µm) and large (>10 µm) vessels per brain section at 450 kPa and 750 kPa 1 hour following 1.5 MHz FUS sonication (N=4 mice / group; **p<0.01; #p = 0.0569; Student’s t-test). (**B**) Quantification of the fraction of small and large leaky vessels at 450 kPa and 750 kPa one hour after FUS sonication. The fraction of leaky vessels is higher for larger, than smaller, vessels at high pressure and there are significantly more large leaky vessels at high than low pressure (N=4 mice/group; *p<0.05; two-way ANOVA with Sidak’s multiple comparison test). (**C**) Quantification of the average number of leaky small (<10 µm) and large (>10 µm) vessels per brain section at 72 hours following 1.5 MHz FUS sonication at 450 kPa and 750 kPa (N=4 mice / group; **p<0.01, ##p = 0.0682; Student’s t-test). (**D**) Quantification of the fraction of small and large leaky vessels at 750 kPa compared to 450 kPa at 72 hours after FUS sonication (N=4 mice/group; ***p<0.001, **p<0.01 two-way ANOVA with Sidak’s multiple comparison test). (**E-H”; K-N”**) Maximum intensity projection images of 25µm-thick caudate putamen sections from *Tg::eGFP-Claudin5*^+/-^ mice sonicated at 450 (**E-F”**) and 750 kPa (**G-H”**). The images show capillaries within the ipsilateral and contralateral caudate / putamen regions 1 (**E-H”**) and 72 (**K-N”**) hours after FUS sonication. eGFP (green) labels TJs between endothelial cells. Biocytin-TMR tracer leakage from blood vessels (red) is seen exclusively in the parenchyma of the treated hemisphere. GLUT-1 (white) labels endothelial cells. The absence of TJ strand in vessels is indicated by white arrowheads. (**I-J**) Quantification of the fraction of TJ strands with gaps between 0.4-2.5 µm (**I**) and > 2.5 µm (**J**) at 450 kPa and 750 kPa. The fraction of TJ strands with gaps > 2.5 µm is significantly different in the ipsilateral compared to the contralateral hemisphere at 750 kPa (N=4 mice / group; *p < 0.05; one-way ANOVA with Sidak’s multiple comparison test). (**O-P**) Quantification of the fraction of TJ strands with gaps between 0.4-2.5 µm (**O**) and > 2.5 µm (**P**) at 450 kPa and 750 kPa. The fraction of TJ strands with gaps greater than 2.5 µm is significantly different in the ipsilateral compared to the contralateral hemisphere at 750 kPa (N=5 mice / group; *p < 0.05; one-way ANOVA with Sidak’s multiple comparison test). All data are presented as mean± standard deviation. Scale bars = 20 µm. See also **Figure S2**.

We analyzed the morphology of eGFP^+^ TJs in *Tg::eGFP-Claudin5^+/-^*mice using confocal microscopy 1 and 72 hours after FUS for both pressures. Since most (∼90%) leaky vessels were small (<10 µm), we analyzed the fraction of capillary TJs containing larger than 2.5 µm in both ipsilateral and contralateral caudate and putamen regions, since TJ gaps correlate with biocytin-TMR leakage across the BBB in animal models for ischemic stroke or multiple sclerosis ^5,39,43^. Within 1 hour post-FUS, biocytin-TMR^+^ leaky capillaries contained eGFP^+^ TJ strands with gaps (>2.5 µm) at both pressures, although these were more pronounced at 750 kPa (**Figure 2E-H”; Figure S3**). There was a 1.5-fold increase in the fraction of TJ strands with gaps >2.5 µm in leaky capillaries for both pressures at 1 hour after FUS, although the lower pressure had a higher variability (**Figure 2I, J**). Moreover, a large number of leaky capillaries lacked completely TJ strands at 750 kPa, whereas these were absent at 450 kPa (**Figure 2H’, F’**). By 72 hours, although there was no difference in the fraction of capillary TJ with gaps >2.5 µm between the ipsilateral and contralateral regions at 450 kPa (**Figure 2K-L”, O**), there was still a significant fraction of capillary TJ with gaps >2.5 µm in the ipsilateral hemisphere at 750 kPa (**Figure 2M-N’, O**). Thus, high acoustic pressures induce persistent structural damage of capillary TJs by confocal microscopy.

Arterioles constitute a small percentage of the leaky vessels at both 1 and 72 hours (**Figure 2A-D**); however, a compromised BBB at the arteriole level can cause significant damage to the brain parenchyma. One hour following FUS, TJ strands with gaps >2.5 µm were found in leaky biocytin-TMR^+^ arterioles at both low and high pressures (**Figure S4A-D”**). However, there was no significant difference in the fraction of TJ strands with gaps >2.5 µm between the ipsilateral and contralateral regions at 450 kPa, although they were higher in the ipsilateral side (**Figure S4A-B”, I, J**). At 750 kPa, the fraction of TJ strands with gaps >2.5 µm or obliterated junctions (loss of entire TJ strands) was 4-fold higher in the ipsilateral compared to contralateral regions (**Figure S4D-D”, I, J**). Thus, high acoustic pressures induce more structural damage to TJs in arterioles that capillaries.

We examined TJ morphology at 72 hours to determine whether junctions were repaired. At 450 kPa, there was no difference in TJ strands with gaps >2.5 µm between the two hemispheres and there was no tracer leakage (**Figure S4E-F”, I, K**). At 750 kPa, the fluorescent tracer could be seen emerging from the vasculature where TJ strands were still damaged (**Figure S4G-H”**). The fraction of TJs with gaps >2.5 μm was 6-fold higher in the arterioles of the ipsilateral compared to the contralateral hemisphere at 72 hours after high pressure FUS (**Figure S4 K**). In summary, transient BBB opening seen at low pressures correlates with transient loss and repair of TJ strands. However, high acoustic pressures induce persistent structural damage to TJs in both capillaries and arterioles that cause long-term BBB damage.

Tight regulation of caveolae-mediated transcytosis ^3–5^ is a key feature of the BBB, prompting us to evaluate whether upregulation in caveolar-mediated transcellular transport could account for the transient BBB opening at low pressure. We compared BBB permeability after FUS between *Tg::eGFP-Claudin5^+/-^* mice and *Tg::eGFP-Claudin5^+/-^*; *Cav1*^-/-^ lacking caveolae (Lutz et al., 2017). There was no difference in BBB opening volume assessed by MRI, and the area or intensity of biocytin-TMR leakage in the caudate and putamen regions was similar between the two genotypes (**Figure S5**). These findings suggest that transient BBB opening at safe pressures is likely not mediated by upregulation in caveolar-mediated transport in CNS ECs, but by transient reorganization of TJs.

### Microglial activation is transient at low, but persistent at high, pressures around leaky CNS vessels

In response to brain injury and BBB damage, microglia change their ramified morphology rapidly and migrate towards the site of injury.^46–48^ We examined microglial responses at both 1 and 72 hours after FUS-mediated BBB opening in the caudate putamen region (**Figure S6**). One hour following FUS-mediated BBB opening, there was no difference in the number of Iba1^+^ cells on the ipsilateral versus contralateral regions at both pressure (**Figure 3A-E**). Ameboid microglia were localized more prominently near the site of BBB leakage at 450 kPa (**Figure 3B-B’”**) and showed thickened processes with 750 kPa (**Figure 3D-D”’**). At low pressure, there was no significant difference in the Iba1^+^ area (microglia or macrophages) between the two hemispheres (**Figure 3F**). In contrast the Iba1^+^ area was increased in the ipsilateral side at high pressure with myeloid cells displaying thickened and elongated processes towards the site of leakage (**Figure 3D-D’”, F**). By 72 hours post FUS, when the BBB is primarily closed at 450 kPa, there was no difference in the number and area covered by Iba1^+^ microglia or macrophages between the two hemispheres which were primarily localized at the sites of BBB leakage (**Figure 3G-H’”, K, L**). At high pressure, there was a 3-fold increase in the number and a 2.8-fold increase in the area of Iba1^+^ cells within the ipsilateral compared to the contralateral region (**Figure 3 K, L**). Microglia or macrophages near the site of BBB leakage showed detracted processes and enlarged cell bodies at 750 kPa (**Figure 3J-J”’**). There was also a 4-fold increase in CD68^+^ area in the ipsilateral compared to the contralateral regions at 750 kPa (**Figure 3M-Q**), consistent with phagocytic myeloid cells.

**Figure 3.**
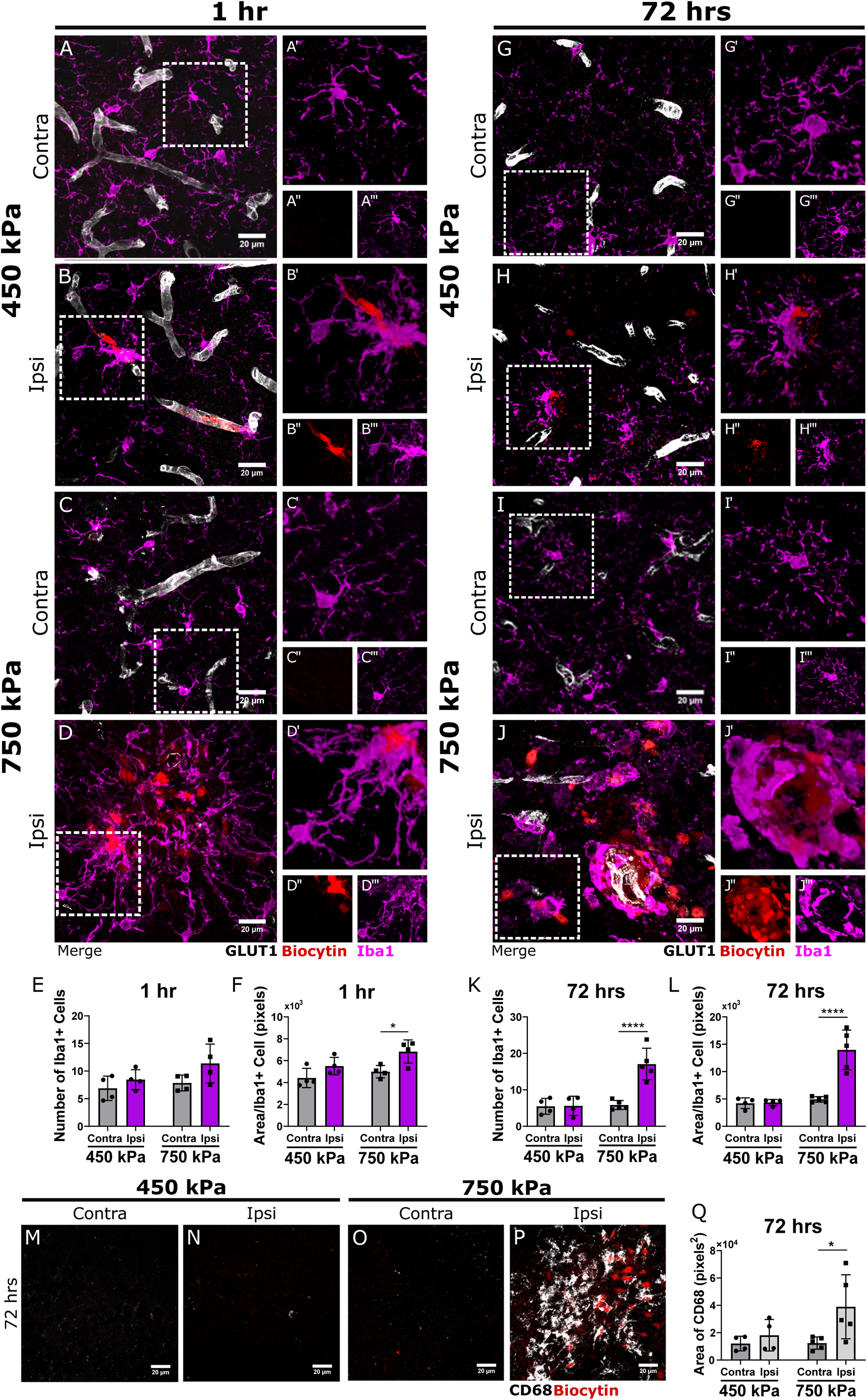
Persistent microglial present only at unsafe 750 kPa pressures after 72 hours following FUS with microbubbles BBB opening. (**A-D’’’; G-J”’**) Maximum intensity projection images (**A”-J”**) and 3D reconstructed volumes (**A”’-J”’**) of 25µm-thick caudate putamen sections from *Tg::eGFP-Claudin5*^+/-^ mice at 1 (**A-D”’**) and 72 (**G-J”’**) hours following 1.5 MHz FUS sonication at 450 kPa (**A-B’’’; G-H”’**) and 750 kPa (**C-D’’’;I-J’”**) pressure. Iba1 (purple) labels microglia or macrophages, biocytin-TMR tracer (red) labels leakage from blood vessels, and GLUT-1 (white) labels blood vessels. (**E, F, K, L**) Quantification of the number of Iba1^+^ cells (**E, K**) and Iba1^+^ cell area (**F, L**) on the contralateral and ipsilateral hemisphere one (**E, F**) and 72 hours (**K, L**) following sonication at 450 kPa (N=4 mice) and 750 kPa (N=5 mice) (*p<0.05, ****p<0.0001; one way ANOVA with Sidak’s multiple comparison test was performed between contralateral and ipsilateral hemispheres). (**M-P**) Maximum intensity projection images showing CD68 positive cells 72 hours following FUS mediated BBB opening at 450 kPa (**M-N**) and 750 kPa (**O-P**). CD68 labels phagocytic microglia or macrophages and biocytin-TMR tracer labels leakage from vessels. (**Q**) Quantification of the CD68^+^ area on the contralateral and ipsilateral hemispheres at 450 kPa (N=4) and 750 kPa (N=5) 72 hours after FUS sonication. (*p<0.05. one-way ANOVA with Sidak’s multiple comparison test). All data are presented as mean ± standard deviation. Scale bar=20 µm. See also **Figure S6**.

### Fibrinogen crosses the BBB following FUS-mediated BBB opening at both pressures and is associated with activated microglia

Fibrinogen is a plasma protein involved in coagulation, inflammation and tissue repair that does not normally cross the BBB.^49^ However, following BBB disruption in neuroinflammation or neurodegeneration, fibrinogen activates microglia initiating neuropathological deficits.^50–54^ We analyzed the area of fibrinogen deposition into the CNS after 1 and 72 h of FUS-mediated BBB disruption at both pressures. One hour following the FUS sonication, fibrinogen was present outside of the vasculature in the ipsilateral hemisphere at both pressures (**Figure S7A-D”**) and the area of fibrinogen leakage was higher on the ipsilateral compared to the contralateral regions at both pressures (**Figure S7E**). Fibrinogen leakage was present only near vessels with biocytin-TMR leakage where there was ameboid microglia (**Figure S7B’’, D’’**). By 72 hours, we could not detect any fibrinogen in the parenchyma at any pressure (**Figure S7F-I, J**). Fibrinogen was contained within the vasculature at 450 kPa and in close association with Iba1^+^ cells at both pressures (**Figure S7G’, G’’, I’, I’’**).

### Astrocytic hypertrophy is present by 72 hours following FUS mediated BBB opening at both pressures

To investigate the astrocytic response following FUS-mediated BBB opening, we quantified the area of GFAP^+^ astrocytes. There was no significant difference in the area of GFAP^+^ astrocytes between ipsilateral and contralateral hemispheres at 1 hour post-FUS for both pressures (**Figure S8A-E**). By 72 hours, there was a significant increase in GFAP^+^ astrocytes in the ipsilateral hemisphere at both pressures, although more prominent for the high pressure (**Figure S8F-J**). Thus, FUS induces changes in astrocyte morphology even at safe pressures, despite restoration of BBB function.

### Single cell RNA-sequencing reveals FUS-induced transcriptional changes related to injury and repair in ECs acutely regardless of the pressure

To determine the transcriptomic response of ECs to low and high pressures at 1 and 72 hours after FUS, we dissociated the caudate and putamen regions from *Tg::eGFP-Claudin5*^+/-^ brains after bilateral FUS into single cells, FACS-sorted eGFP^+^ ECs and performed single-cell RNA-sequencing (scRNAseq) and analysis as described^55^ (**Figure 4A, Table S1**). After quality filtering^56^ and *in silico* EC selection based on defined markers [*Pecam1*, *Cldn5*, and *Erg*],^57^ we performed graph-based clustering to separate ECs sequenced from 10 brain samples (2 samples per condition and 2 healthy controls; **Table S1**) into distinct clusters based on transcriptomic expression profiles using high resolution. This approach separated ECs into 20 clusters visualized using a Uniform Manifold Approximation and Projection (UMAP) representation (**Figure 4B-D)**. While these EC clusters are not necessarily biologically distinct, this approach (over-clustering and re-merging based on computational or biological meaning) avoids pitfalls arising from incorrect clustering boundaries using lower resolution.^58^ We assigned a subtype identity to each EC cluster based on expression of canonical markers for arteries, capillaries and venules (**Figure 4E, F**). Expression of arterial and venous markers peaked at opposite ends of the UMAP plot separated by capillary clusters, suggesting that arteriovenous (AV) zonation was a major variation axis in the EC data (**Figure 4E, F**). The second largest variation axis was the FUS pressure at 72 hours versus control conditions (healthy ECs). Capillary ECs were the largest population, arterioles ECs comprised the second one, and venule ECs were the smallest population for each condition, as expected based on their distribution in the striatum and putamen (**Figure 4G**).

**Figure 4.**
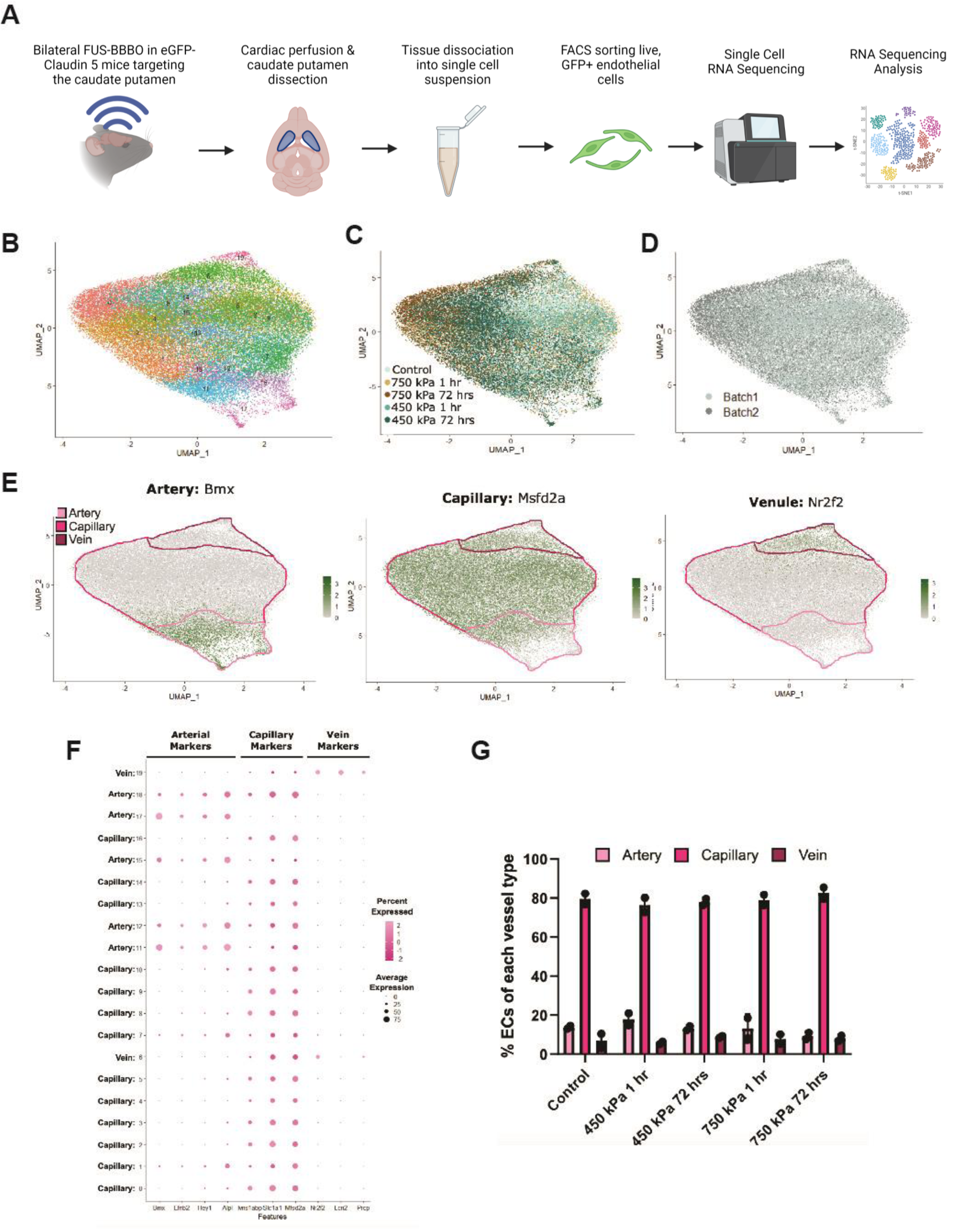
Single-cell RNA sequencing of endothelial cells after FUS and identification of EC subtypes. (**A**) Experimental workflow for single-cell RNA sequencing experiments of ECs after FUS. (**B-D**) Uniform manifold approximation and projection (UMAP) of ECs integrated and color coded by Seurat cluster, treatment group, and sequencing batch. (**E**) Clusters of arterial, capillary and venous cells identified by expression of canonical markers Bmx, Msfd2a and Nr2f2 respectively in the UMPA plot. (**F**) Dotted plot of canonical markers used to identify clusters to be classified as arterial, capillary and venous cells. The size of the dot indicates the number of cells and the color the intensity of expression. (**G**) Dotted bar plot showing the percentage of arterial, capillary, and venous ECs for each treatment group. See also **Figure S10** and **Tables S1-S4**.

To identify key transcriptomic signatures differentiating ECs from each condition, we performed differential gene expression analysis comparing either FUS 1 hour (there were no differences between 450kPa and 750 kPa), FUS 450 kPa 72 hours, or FUS 750kPa 72 hours to control-enriched EC clusters for each vessel subtype (**Tables S2-S4**). The short-term transcriptomic responses of capillary ECs to 450 and 750 kPa 1 hour post-FUS were statistically indistinguishable, so they were grouped together for further analysis. This data was used to conduct gene ontology (GO) analysis in DAVID and gene set enrichment analysis (GSEA) using curated and comprehensive gene catalogs for various cellular and molecular pathways and functions from the Molecular Signatures Database v6.1 (*MSigDB:* software.broadinstitute.org*/ gsea/msigdb/genesets.jsp*),^59,60^ and determine which biological processes were significantly up- and downregulated for each condition (**Tables S5-S8**). The transcriptomic analysis of capillary ECs one hour-post-FUS revealed upregulation of several genes and GO terms related to “cell death (e.g. *Txnip*)”, “intracellular signal transduction (e.g. *Hes1, Smad1, Smad6*),” “regulation of cell proliferation (e.g. *Cdhkn1a*),” and “vasculature development (e.g. *Klf4, Hes1*)” relative to untreated controls (**Figure 5A, D; Tables S2, S5**). However, the acute capillary ECs response to FUS was more complex as other transcripts related to “vascular development” (*e.g. Pdgfb, Kdr, Lef1, Nrp1*) were downregulated compared to control ECs (**Figure 5A, D, Tables S2, S5**). This complex transcriptional EC response suggests initiation of a potential repair response in ECs in additional to the injury response in the acute phase (1 hour) post-FUS.

**Figure 5.**
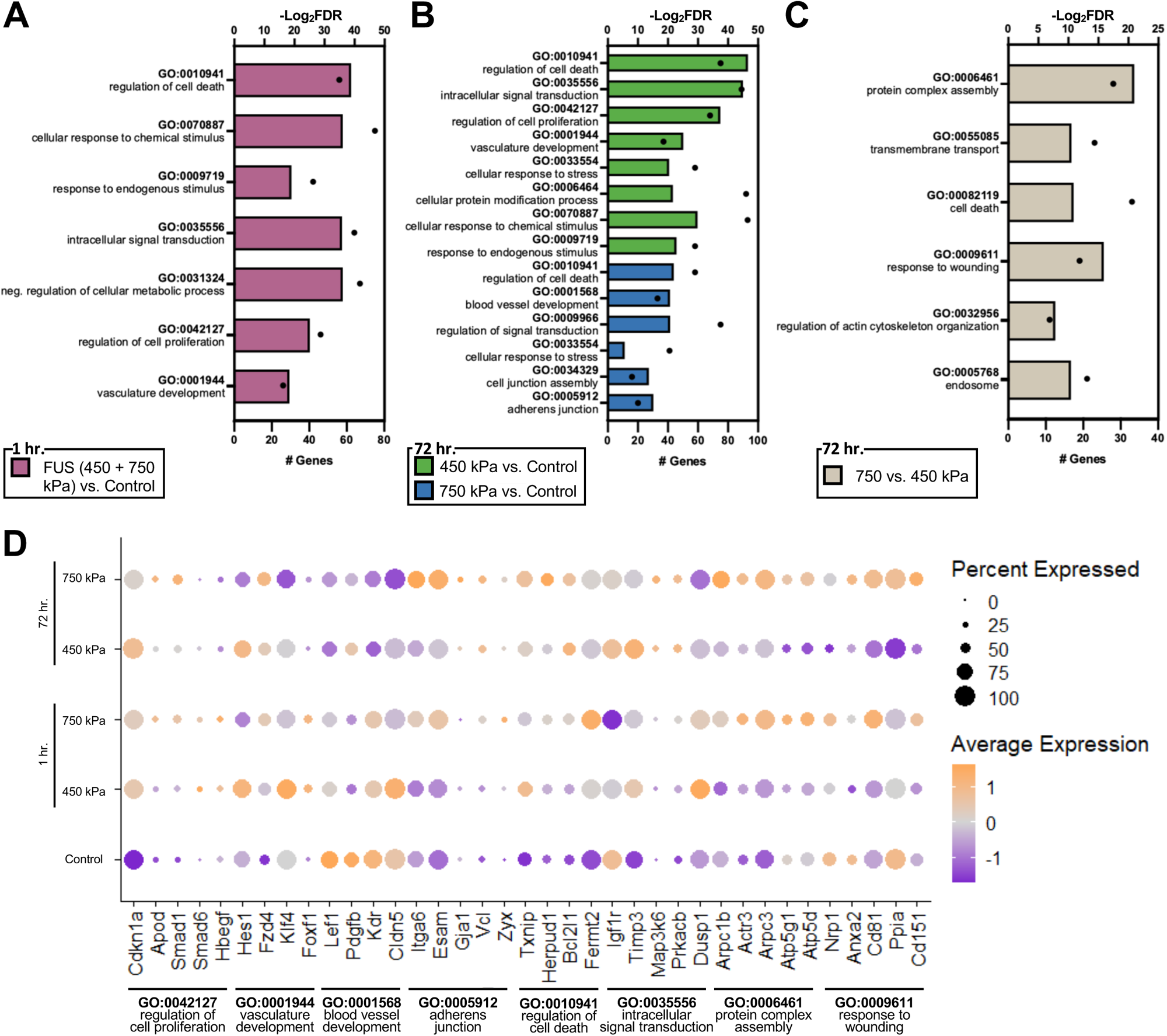
Gene ontology analysis of differentially expressed genes in capillary ECs for each condition from single cell RNA sequencing. (**A**) Gene ontology terms upregulated in FUS (450 + 750 kPa) capillary cells compared to untreated controls 1 hour post-FUS-BBBO. (**B**) Gene ontology terms upregulated in capillary cells exposed to 450 kPa (green) and 750 kPa (blue) compared to untreated controls, 72 hours post-FUS-BBBO. (**C**) Gene ontology terms upregulated in capillary cells exposed to 750 kPa compared to 450 kPa cells 72 hours post-FUS-BBBO. In all bar plots the bar represents the -Log_2_(FDR) (top axis), and the dot represents the number of significantly annotated genes (bottom axis). (**D**) Dot plot showing relative expression of key genes annotated to the upregulated gene ontology terms for each treatment group and untreated controls. The size of the dot indicates the number of cells and the color the intensity of expression. See also **Tables S5-S7**.

GSEA analysis of capillary ECs revealed upregulation of “inflammation”, “notch signaling” and “endothelial to mesenchymal transition” at 1 hour post-FUS (**Figure S9A**). A similar transcriptional response was observed also in arterioles and venules 1 hour post-FUS. Arterioles exposed to 750 kPa showed increased expression of transcripts annotated to “regulation of cellular component movement” and “cell-substrate junction” pathways, and venules exposed to 450 kPa demonstrated upregulation of “blood vessel development” and “regulation of cellular component movement” (**Figure S10A-B, Table S2, S6, S7**). Therefore, all three EC subtypes respond dynamically at the acute phase (1 hour) post FUS to upregulate transcripts related to both injury processes (e.g. “cell death”, “inflammation”) and repair processes (e.g. “cell proliferation”, “cell migration”, “vascular development”) regardless of the applied pressure.

### Transcriptomic analysis after 72 hours reveals that ECs upregulate processes related to angiogenesis and barriergenesis in particular at high pressures

At 72 hours post-FUS the BBB is restored in mice exposed to low pressure (450 kPa), whereas it remains open in mice exposed to high pressure (750 kPa; **Figure 1**). This was reflected in distinct transcriptomic EC responses between the two pressures by 72 hours. GO term analysis of DEGs upregulated in capillary ECs at 72 hours post - 450 kPa compared to control ECs revealed upregulation of pathways such as “regulation of cell death” (e.g. *Txnip, Bcl21b*), “intracellular signal transduction,” “regulation of cell proliferation” (e.g. *Cdnk1a*), “vascular development” (*Notch1, Notch4, Hes1, Klf4, Clic4*) and “cellular response to stress” (e.g. *Slc2a1, Slc38a2*) (**Figure 5B, D, Table S3, S5**). Surprisingly, GSEA analysis of DEGs at 72 hours versus 1 hour post-450 kPa or control ECs showed downregulated pathways for “blood-brain barrier”, “cell proliferation”, “angiogenesis”, “EC migration”, “tip cells”, “Wnt/β-catenin, Notch and TGF-β signaling” (**Figure S9A, B, Table S8**). These data indicate that CNS ECs are likely still undergoing vascular growth and BBB maturation at 72 hours post-FUS at low pressures.

Similarly, the GO term analysis of DEGs upregulated in capillary ECs at 72 hours post-750 kPa showed also upregulation of pathways such as “regulation of cell death” (e.g. *Txnip, Bcl21b*), “intracellular signal transduction,” “regulation of cell proliferation” (e.g. *Cdnk1a*), and “vascular development” (*e.g. Notch1, Notch4, Hes1, Klf4, Clic4*) compared to control ECs (**Figure 5B, D**). In contrast to 450 kPa, GSEA analysis revealed that processes such as “blood-brain barrier”, “inflammation”, “tip cells” were upregulated at 72 hour post-750 kPa, whereas those to “angiogenesis”, “EC migration”, VEGF, TGF-β and non-canonical Wnt signaling” were downregulated compared to controls (**Figure S9A, B, Table S8**). Pathway analysis of upregulated DEGs in ECs after 72 hours after 750 kPa compared to 450 kPa revealed the presence of pathways related to “protein complex assembly,” (e.g. *Arpc1b, Arpc3, Actr3, Atp5d, Atp5g1*), “regulation of cell death,” and “response to wounding” (e.g. *CD81, CD151, Ppia*) (**Figure 5C, D**). These transcriptome findings are consistent with the concept that ECs exposed to 450 kPa mostly recovered by 72 hours post-FUS, whereas those exposed to 750 kPa are still undergoing cell death and early repair processes, since they were destroyed by the high pressure.

### Several major signaling pathways related to angiogenesis and barriergenesis change dynamically in ECs after FUS

In order to understand which signaling pathways critical for angiogenesis and barriergenesis change differentially in ECs after low and high FUS pressure, we performed a GSEA analysis for specific pathways on our scRNAseq data. TGF-β signaling, which functions to inhibit cell proliferation, suppress inflammation and promote vascular development, is upregulated in capillary ECs one hour post-FUS consistent with an increase in inflammatory responses (**Figure 6A; Tables S5, S8).** However, this pathway is largely downregulated by 72 hours relative to either control ECs, one hour post-FUS at 750 kPa, or 72 hours post-FUS 450 kPa (**Figure 6A; Tables S5, S8**), consistent with reduced inflammation and promotion of angiogenesis. Several TGF-β signaling genes, Tgfb2 and Tgfbr2, Smad 1 and Smad 6, were significantly downregulated in the 750 kPa group 72 hours post-FUS relative to other conditions (**Figures 5D, 9B**).

**Figure 6.**
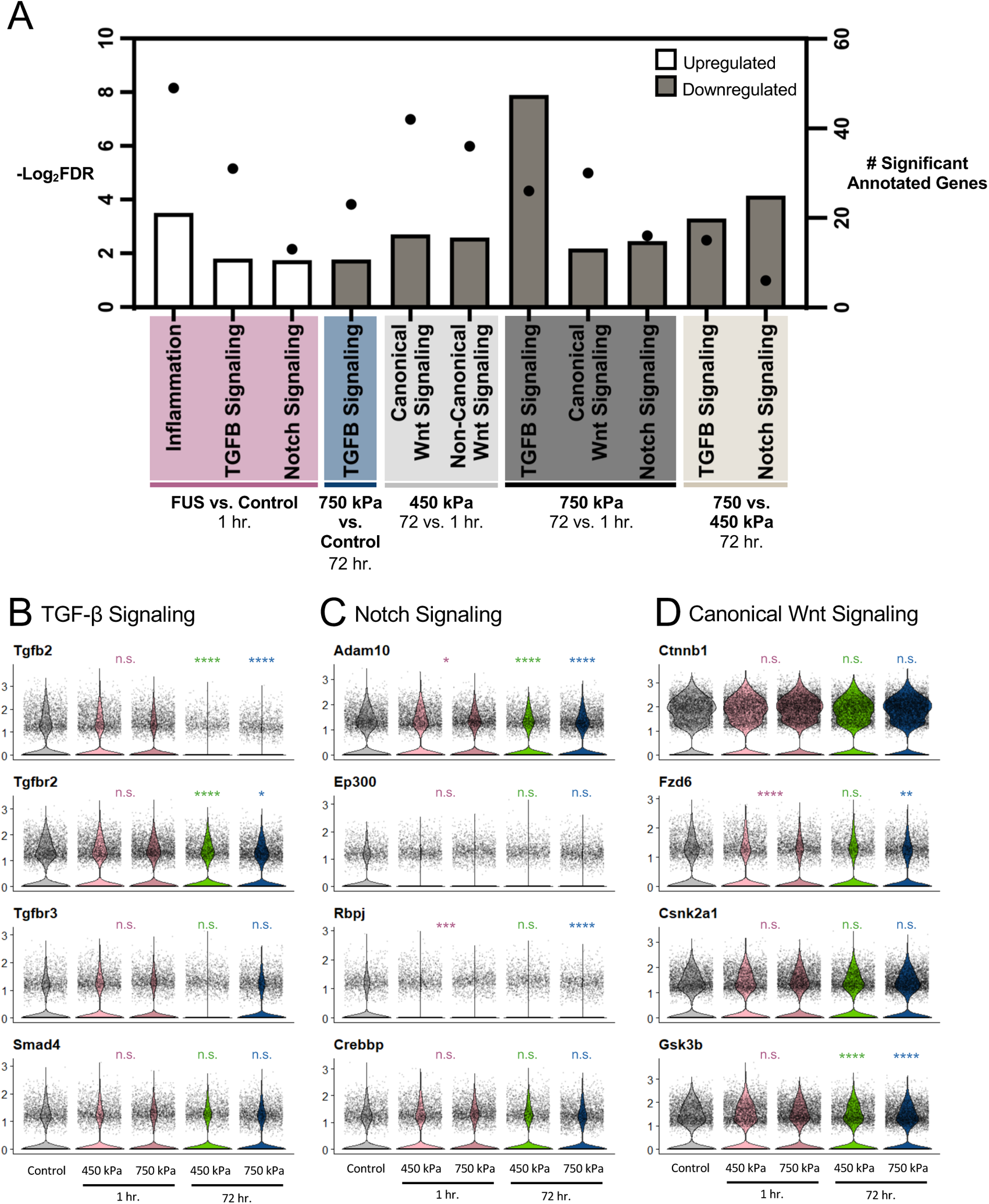
Gene set enrichment analysis (GSEA) results for capillary cells. (**A**) GSEA terms up and downregulated in capillary cells for the indicated comparisons (P value cutoff < 0.3). Bars represent the -Log_2_(FDR) values (left y-axis), and dots represent the number of significant genes annotated to each term (right y-axis). Comparison label convention is “Measured vs. Reference”. Namely, “FUS vs Control” represents changes observed in FUS groups using the Control group as reference. (**B-D**) Violin plots showing the expression of representative genes annotated to TGF-β Signaling, Notch Signaling, and Canonical Wnt Signaling. Statistical significance was measured between each group and the Control using pairwise Wilcoxon rank sum tests to determine the adjusted p-value. FUS groups 1 hour post-FUS-BBBO were pooled for this analysis (450 + 750 kPa). Significance values are indicated as follows: * P ≤ 0.05, ** P ≤ 0.01, ***

Notch signaling, which is an important regulator of angiogenesis and EC proliferation, was a second signaling pathway that changed dynamically in ECs after FUS. Notch signaling components (e.g. (*e.g. Notch1, Notch4, Hes1, Rbpj, Adam100)* were upregulated significantly in capillary ECs one hour post-FUS compared to control ECs (**Figures 5D, 6B; Tables S5, S8**). However, the Notch pathway was largely downregulated by 72 hours post-FUS in capillary ECs, relative to those isolated from one hour post-FUS at 750 kPa, or 72 hours post-FUS at 450 kPa (**Figures 5D, 6B; Tables S5, S8**). This trend was also consistent for arteriole ECs. In addition, arteriole ECs also showed a decrease in VEGF signaling at 72 hours post 750 kPa relative to untreated controls (**Figure 6A**). However, when comparing arterioles exposed to 750 kPa to those exposed to 450 kPa 72 hours post-FUS, GSEA analysis reveals increased expression of TGF-β and VEGF signaling. Therefore, several signaling pathways critical for angiogenesis and EC proliferation change dynamically after FUS, but they remain lower in high pressure at long-term time points consistent with more blood vessel damage.

Wnt/β-catenin signaling is critical for CNS angiogenesis and BBB maturation, prompting us to examine its EC transcriptional changes after FUS carefully. GO and GSEA analyses did not show any significant differences in genes annotated to Wnt/β-catenin signaling in either low or high pressure at the acute phase, compared to the untreated control group. However, canonical Wnt signaling was decreased at both pressure groups at 72 hours compared to one hour post-FUS (**Figure 6A, Tables S5, S8**). Additionally, arterioles exposed to both pressures showed decreased expression of Wnt/β-catenin signaling at 72 compared to one hour post-FUS (**Table S6**). Overall, canonical Wnt signaling is higher at the acute than the sub-acute (72 hours) phase post-FUS for both pressures and at lower than higher pressures by 72 hours consistent with continuous “EC proliferation” and “angiogenesis” at this time point at 750 kPa. Overall, this analysis demonstrates a reduction in the injury response and “return to normalization” at both pressures by 72 hours relative to one hour post-FUS, since Wnt/β-catenin signaling is reduced once angiogenesis and BBB maturation are complete.^1,2^

We validated some of the scRNA-seq findings related to “angiogenesis” using a fluorescent in situ hybridization (FISH) assay for two angiogenesis markers *Egfl7* and *Angpt2* ^55^ (**Figures 7, S11**) in combination with immunofluorescence for a vessel marker Glut-1 (Slc2a1). Violin plots from scRNA-seq data revealed significant increase for *Egfl7* and *Angpt2* mRNAs in the pooled 1 hour pressure group, and at 72 hours in both 450 and 750 kPa relative to untreated controls as determined by Wilcoxon rank sum test (**Figures 7A, S11A**). By FISH, *Egfl7* mRNA was slightly upregulated in Glut1-positive vessels exposed to 450 kPa on the ipsilateral compared to contralateral hemispheres, whereas we could not detect any *Angpt2* mRNA upregulation (**Figure 7B&C**). In contrast both *Egfl7* and *Angpt2* mRNAs were highly upregulated in Glut-1+ vessels exposed to 750 kPa on the ipsilateral compared to contralateral hemisphere (**Figure S11B&C**). These findings validate the scRNAseq data and the GO and GSEA analyses which indicate “angiogenesis” at 72 hours in both high and low pressure conditions, with a more extreme and sustained response in the 750 kPa compared to 450 kPa group. In conclusion, the BBB defects due to FUS promote an “angiogenic” program in CNS blood vessels likely to drive vascular repair.

**Figure 7.**
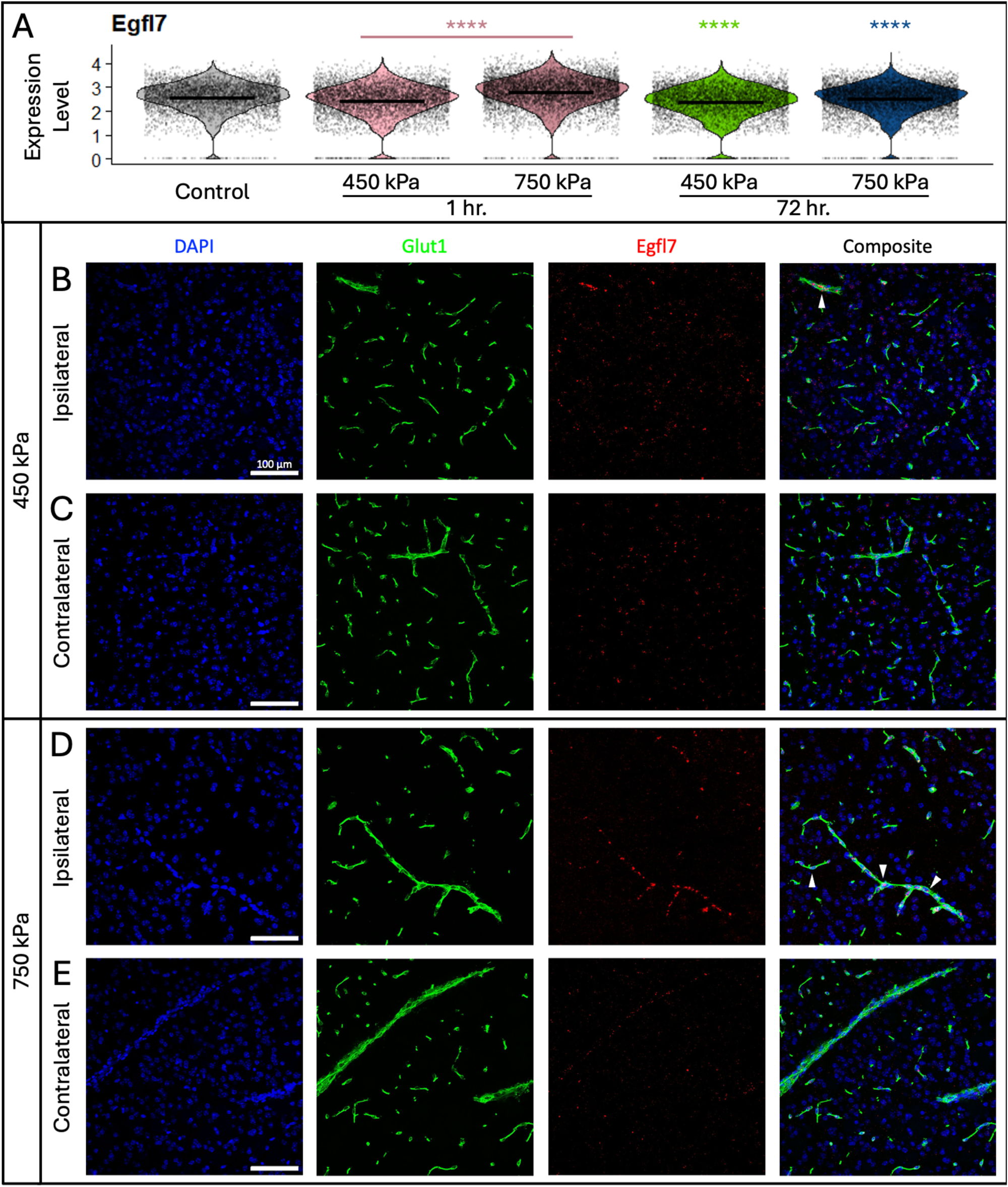
Glut1-positive endothelial cells exposed to 750 kPa exhibit a sustained, FUS-induced increase in Egfl7 expression by fluorescence in situ hybridization assay. (A) Violin plots showing expression of Egfl7 across each of the treatment groups. Mean expression level is indicated by the bar in each plot. Statistically significant differences between each group and untreated controls are determined by Wilcoxon rank sum test. **** P ≤ 0.0001. EC’s exposed to 450 kPa (B) exhibit elevated levels of Egfl7 expression compared to the untreated contralateral hemisphere (C). EC’s exposed to 750 kPa (D) exhibit markedly greater expression of Egfl7 relative to the untreated contralateral hemisphere (E). White arrowheads indicate Glut1-positive vessels with positive Egfl7 expression.

## MATERIALS AND METHODS

### Animals

All experimental procedures involving animals were approved by the Columbia University Institutional Animal Care and Use Committee (IACUC) and in accordance to the Office of Laboratory Animal Welfare and the Association for Assessment and Accreditation of Laboratory Care regulations. A total of eighteen (N=18) male and female C57BL6 or *Tg::eGFP-Claudin5* and three (N=3) male *Tg::eGFP-Claudin5^+/-^ Cav1^-/-^* 12-14 weeks old mice (20-25 g) were used for this study. Animals were housed under standard conditions (12 h light/dark cycles, 22°C), were fed a standard rodent chow (3kcal/g; Harlen Laboratories, Indianapolis, IN, USA) and drank distilled water. All mice had access to their diets ad libitum. Animals were group housed and were randomly selected into stable (Pressure=0.45 MPa) or inertial cavitation (Pressure=0.75 MPa) groups.

### Microbubble Preparation

In-house lipid-shelled microbubbles were manufactured as previously published.^61^ Briefly, the 1,2-distearoyl-sn-glycero-3-phosphocholine (DSPC) and polyethylene Glycol 2000 (PEG2000) were mixed at a 9:1 ratio. Two milligrams of the mixture was dissolved in a 2-ml solution of filtered PBS/glycerol (10% volume)/propylene glycol (10% volume) with a sonicator (Model 1510, Branson Ultrasonics, Danbury, CT, USA) and stored in a 5 ml vial. The remainder of the vial was filled with decafluorobutane (C_4_F_10_) gas. The vial was then activated via mechanical agitation using VialMix^TM^ shaker (Lantheus Medical Imaging, N. Billerica, MA) for 45 s. The formed microbubbles were analyzed with a Coulter Counter Multisizer (Beckman Coulter Inc., Fullerton, CA).

### FUS Blood-Brain Barrier Opening

A single-element 1.5 MHz center frequency focused ultrasound transducer (center frequency: 1.5 MHz, focal depth: 60 mm, radius: 30 mm, axial full width half-maximum intensity: 7.5 mm, lateral full-width half-maximum intensity: 1 mm, Imasonic, France) was used for all BBB openings in this study. To measure the beam profile, the transducer was calibrated using a needle hydrophone (HGL-0400, Onda Corp., Sunnyvale, CA) in a tank of deionized, degassed water. The FUS transducer was driven by a function generator (Agilent, Palo Alto, CA, USA) though a 50-dB power amplifier (E&I, Rochester, NY, USA). A pulse-echo transducer (center frequency: 7.5 MHz, focal depth: 60 mm, radius 13 mm; Olympus NDT, Waltham, MA) was confocally aligned with the FUS transducer to monitor cavitation events. The pulse-echo transducer was driven by a pulse-receiver (Olympus NDT, Waltham, MA) connected to a digitizer (Gage Applied technologies, Inc., Lachine, QC, Canada) for data acquisition. During passive cavitation detection (PCD) acquisition the pulse-receiver operated in “receive mode” and served as an amplifier during PCD acquisition. The transducer setup was then mounted onto a three-dimensional positioning system (Velmex Inc., Lachine, QC, Canada) for accurate targeting.

A bolus of 1 µL/g of body mass of polydisperse microbubbles diluted in (8X10^8^/mL, mean diameter: 4.5 µm) with 100 µL was intravenously injected immediately preceding the sonication. For experiments involving *ex vivo* visualization of biocytin diffusion, mice survived for 1 hr were co-injected with the fluorescent tracer biocytin-tetramethylrhodamine (870 Da, Life Technologies). The caudate putamen is a biologically relevant structure for diseases such as Alzheimer’s and Parkinson’s, and a structure that has been previously targeted for drug delivery^62^. Targeting of the caudate putamen was achieved by first locating the lambda structure though an intact dilapidated scalp and skull as previously published ^63^. Briefly, a water cone filled with deionized (DI), and degassed water was fixed to the FUS transducer, which was submerged in a water bath of DI and degassed water, and finally coupled to the shaved scalp using degassed ultrasound gel. A metallic wire with orthogonal parts was placed above the lambda structure and a C-mode ultrasound was performed to locate the cross and below it, the lambda structure. The FUS transducer was then moved via the positioner to the following coordinates from lambda: AP 4.5 mm, ML 2.5 mm and DV -3.5 mm for caudate targeting and a single sonication was performed. The FUS parameters used were as follows: frequency 1.5 MHz, peak-rarefactional pressure 0.45 MPa or 0.75 MPa, pulse length 10 ms, pulse repetition frequency 10 Hz, and for a duration of 120 s.

### Magnetic Resonance Imaging

One hour following the ultrasound procedure, all animals underwent scanning with the 9.4 T MRI system (Bruker Medical, Boston, MA). The mice were placed in a birdcage coil (diameter 35 mm), while being anesthetized with 1 -2% isoflurane and respiration rate was monitored throughout the imaging sessions. MR images were acquired using a contrast-enhanced T1-weighted 2D FLASH sequence (TR/TE 230/3.3 ms, flip angle: 70°, number of excitations: 6, field of view: 25.6 mm × 25.6 mm, resolution 100 μm x 100 μm x 400 μm), 45 min following the intraperitoneal bolus injection of 0.3 ml gadodiamide (GD-DTPA) (OmniscanTM, GE Healthcare, Princeton, NJ). As previously reported, gadodiamide provides spatial information of the BBB opening, as it will diffuse into the brain where the BBB disrupted, enhancing MR signal temporarily and enabling *in vivo* visualization of the BBB opening volume.^64^ Mice survived for 72 hours, underwent an additional scan of the same sequence, 71 hrs following FUS.

MRI BBB Opening Volume Quantification: Contrast-enhanced T1-weighted MRI images were processed in MATLAB (2019a Mathworks, Natick, MA,USA). Analysis was performed on 15 consecutive coronal images that were manually segmented to remove the skull from the brain. The volume of BBBO, or BBB disruption is defined for the purposes of this study as the volume of the brain into which Gd contrast agent is able to diffuse and be visualized on T1-weighted MRI. This affected volume will be the focus of subsequent histological and transcriptomic analyses. Images were thresholded to generate a binary mask based on the mean plus two standard deviations pixel intensity of an ROI chosen on the contralateral hemisphere. The area of each mask was calculated and summed across all 15 slices to give the final BBB opening volume.

### Immunofluorescence Staining and Imaging

For the biocytin-TMR quantification, mice survived for 72 hours, were injected with biocytin-TMR one hour prior to sacrifice. Either 1 or 72 hours post-sonication, mice were transcardially perfused with 30 mL PBS followed by 60 mL 4% paraformaldehyde. The skull was removed and the brain was soaked in paraformaldehyde for six hours, transferred to PBS overnight, cryoprotected in 30% sucrose for 48 hours, and then frozen on dry ice. Brains were coronally sectioned at 25 µm throughout the caudate putamen region. A series of 4-6 sections within the focus (1 mm) centered at Bregma 0.3 mm were serially selected for each mouse for staining. Tissues were stained for eGFP (1:1000; Sigma-Aldrich, MO), Glut-1 (1:1000; EMD Millipore, MA), DyLight 649 Labeled Griffonia (Bandeiraea) simplicifolia lectin I (1:250; Vector Laboratories, CA), Streptavidin-Alexa594 (1:1000; ThermoFisher) was used to visualize biocytin-TMR distribution in tissues,^39^ Iba-1 (1:100; Abcam), CD68 (1:100; Abcam), Fibrinogen (1:500; EMD Millipore, MA), and GFAP (1:1000, Abcam). Whole brain images were captured with an upright Zeiss AxioImager. Confocal images for analysis of tight junction morphology were captured on a Zeiss LSM700 confocal microscope. For each section, 4-6 images were captured on both the contralateral and ipsilateral hemisphere exclusively within the caudate putamen. Images on the ipsilateral hemisphere were acquired near sites of BBB opening, determined by the presence of Biocytin-TMR. Images were minimally processed with Fiji/ImageJ software (NIH) to enhance brightness and contrast.

### In Situ Hybridization

Fluorescent *in situ* hybridization combined with IF of fresh-frozen sections using antisense mRNA probes for *Egfl7* and *Angpt2* transcripts and Glut-1 (vessel marker) were performed as previously described.^55^ The sections were imaged at 20x using a Zeiss LSM900 confocal microscope.

### Acoustic Signal Analysis

For each FUS pulse transmitted, microbubble cavitation was acquired by a single element PCD, transferred to a digitizer (GAGE Applied Technologies Inc, Lachine QC, Canada), and processed in MATLAB (2017a Mathworks, Natick, MA, USA). Each PCD pulse was transformed into a power spectrum using a fast Fourier transform (fs= 50 MHz) and the resulting energy spectral density was bandpass filtered (3 MHz -9 MHz). The peak value within a 20 kHz bandwidth around each harmonic (nf_c_, n=1, 2… 6, f_c_=1.5 MHz) and ultraharmonic (nf_c_/2, n=3, 5, 7, 9) was found. The cavitation dose is computed as the root mean square (rms) of the spectral amplitude for each time point. SCDu and SCDh are given by the CD of their respective bins. The inertial cavitation dose (ICD) is defined as any CD energy not contributing to SCDh or SCDu. The cumulative cavitation dose is given integrating the amplitude over the entire sonication duration. The cumulative cavitation emissions from microbubbles were normalized to baseline cavitation dose measured prior to IV injection of bubbles.

### Single-Cell RNA Sequencing Preparation

At either 1 or 72 hr. post-FUS, eGFP-Cldn5 mice were deeply anesthetized with 4% isoflurane and perfused for 4 minutes with 4°C, sterile PBS. The brain was dissected into Earl’s balanced salt solution (EBSS) and dissociated into a single cell suspension using an established protocol.^55^ The single-cell suspension was resuspended in 200 μl of CD16/CD32 Fc block (BD Biosciences, Cat. # 553141; 1/200 in FACS buffer) and incubated at room temperature for 15 minutes. Samples were washed with 2 ml of FACS buffer, and 100 μl were kept aside for unstained and fluorescence minus one (FMO) controls. The cells were resuspended in 200 μl of anti-CD31-APC (Biolegend, Cat. # 102410; 1/200 in FACS buffer) antibody to label endothelial cells (ECs) and incubated on ice in the dark for 30-60 mins. Following antibody labeling, the samples were washed twice with 2 ml of FACS buffer and resuspended in 400 μl of propidium iodide (Thermofisher, Cat. # P1304MP; 1/10,000 in FACS buffer). Gates for the surface stain were set using the unstained and FMO controls. The double, eGFP-Cldn5 and CD31-APC positive ECs were sorted out using the BD FACS Aria Flow Cytometer (Columbia Stem Cell Initiative Flow Cytometry Core) and processed for scRNAseq.

## QUANTIFICATION AND STATISTICAL ANALYSIS

### BBB Leakage and Vessel Size Quantification

Whole brain slices were used to quantify the intensity of the fluorescent tracer Biocytin-TMR and the vasculature by immunofluorescence staining for GLUT-1. Leaky vessels were defined as segmented vessels that colocalized with areas of leakage determined by a fixed threshold. Biocytin-TMR leakage was quantified with a custom algorithm in MATLAB (R2017a, Mathworks, Inc., Natick, MA, USA). For quantification of fluorescence intensity, an ROI of the entire sonicated hemisphere containing the caudate putamen and the other brain structures affected by the FUS focus, such as the somatosensory cortex, was selected for each section in the ipsilateral and contralateral side and fluorescence intensity was calculated by subtracting the selected area times the mean fluorescence of the background from the sum of the pixel values of the ROI. Vessels were identified by GLUT-1 staining, and a custom script written in MATLAB (R2017a, Mathworks, Inc., Natick, MA, USA) segmented vessels and measured the diameter. Leaky vessels were defined as segmented vessels that colocalized with areas of biocytin-TMR leakage determined by a fixed threshold. For quantification of the percentage of leaky vessels for small and large caliber vessels, the number of leaky vessel for either small or large vessels was divided by the total number of small or large vessels in the ROI. For each identified vessel the area of biocytin-TMR leakage above a fixed threshold and bioctin-TMR intensity was computed within an ROI centered around the vessel.

### TJ Quantification

Maximum intensity projections were created in order to quantify TJ abnormalities. The number of gaps was quantified for each TJ junction strand in leaking vessels on the ipsilateral side, and control vessels on the contralateral side. Vessel boundaries were determined by GLUT-1 staining. Arterioles, venules, and capillaries were identified based on 1) diameter of the vessel to separate capillaries from arteiroles and venules and 2) tight junction structure to distinguish between the three vessel types. Colocalization with leakage of the fluorescent tracer Biocytin-TMR (870 Da) with venules, identified by their TJ morphology, were not found, therefore analysis of venules was excluded from this analysis. Gaps and protrusions are defined as described.^5,39^ Only TJ strands contained within vessel boundaries were included in the analysis. Tight junction gap length was measured with Fiji/Image J software (NIH) using maximum intensity projection images from Tg::eGFP-Claudin5 mice one hour after FUS. Histograms of the length of tight junction disruption pooled across mice and vessel type demonstrated a bimodal distribution, and a cutoff threshold of 2.5 μm was chosen. Data was sorted based on gap lengths from 0.4-2.5μm and gaps greater than 2.5 μm.

### Immunofluorescence Area Quantification

Maximum intensity projections were created and pixels were thresholded determined by the mean plus two standard deviations of a manually selected ROI containing background noise. For a given immunofluorescent stain, all images underwent the same processing pipeline regardless of hemisphere, pressure, or time point.

### Iba1^+^ Cell Count and Iba1+ Cell Area Quantification

A structural algorithm in MATLAB (R2017a, Mathworks, Inc., Natick, MA, USA) was developed to quantify the number of Iba1 cells. Briefly, maximum intensity projections were created and thresholded for Iba1 signal. Morphological opening and closing was implemented to resolve microglia with cell bodies from stray microglial processes emanating from out of plane microglia. Finally, a Moore-Neighbor tracing algorithm^65^ was implemented to identify individual cells. Pixel area corresponding to each microglial cell body and connected processes was computed to get the Area/Iba1+ cell.

### Iba1 3D volume renderings

3D texture-based volume renderings were generated in Fiji/ImageJ software (NIH) with a resampling factor of 1 using 25 μm thick sections with Iba1 (magenta) and Biocytin-TMR (red) channels.

### Single-Cell RNA Sequencing Analysis

For single-cell RNA sequencing (scRNAseq), the 10x Genomics Chromium platform was used to generate low-depth data (∼2000 genes/cell), which was processed using the Cell Ranger analysis pipeline to align reads and generate feature-barcode matrices. The Seurat R package^56^ was implemented to read the output of the Cell Ranger pipeline and merge cells from all the samples (untreated controls, high and low FUS pressures, 1 hr. and 72 hr. timepoints) into a single R object. The standard pre-processing workflow for scRNAseq data was carried out in Seurat: cells were filtered based on quality control metrics (number of unique genes detected in each cell > 200, total number of molecules detected within a cell > 1,000 and < 50,000, percentage of reads within a cell that map to the mitochondrial genome < 20, etc.), the data was normalized and scaled, and highly variable features were detected. Next, linear dimensionality reduction (PCA) was performed on the scaled data using the previously determined highly variable features, an alternative heuristic method (‘Elbow plot’) was implemented to determine the ‘dimensionality’ of the dataset, and cells were clustered by applying the weighted shared nearest-neighbor graph-based clustering method.^66^ Finally, the PCA dimensions were further reduced into two-dimensional space using the non-linear dimensional reduction technique, t-distributed stochastic neighbor embedding (t-SNE), to visualize and explore the scRNAseq dataset and each of its unique clusters. To ensure the purity of endothelial cells in subsequent analysis, all sequenced cells were clustered and the expression levels of canonical cell-type markers Cldn5, Ptprc, Gfap, Pdgfrb, Map2, and Acta2 were evaluated across the clusters. A cluster of contaminating pericytes was identified and excluded based on high expression of Pdgfrb and Acta2. The remaining, pure endothelial cells were then segmented out and clustered again. The expression of vessel subtype markers such as arterial (Bmx, Efnb2, Hey1, and Alpl), capillary (Ivns1abp, Slc1a1, Mfsd2a) and vein (Nr2f2, Len2, and Prcp) were used to identified arterial, capillary and vein endothelial cells as shown in Figure 2C.

### Differential Gene Expression Analysis

The Seurat “FindMarkers” function with the default Wilcoxon Rank Sum test was implemented to compile the lists of differentially expressed genes, by comparing expression profiles of FUS-BBBO EC clusters to control EC clusters at 450 and 750 kPa, at both acute (1 hr.) and long-term (72 hr.) timepoints.

### Gene Set Enrichment Analysis (GSEA)

Differential gene expression (DE) analysis was performed to compare gene expression variation between experimental groups^67^ and highlight which clusters have further heterogeneity associated with each condition. Following DE analysis, a score was calculated for each gene using the formula: Gene score = -log10(pval) x sign (log2fc). Up- and downregulated genes had a positive and negative score, respectively. The numeric value of the gene score was inversely correlated with the degree of significance, that is, genes that had the highest statistically significant difference in expression and the lowest p-values had the highest score, while non-significant genes with high p-values had a correspondingly low gene score. This score was used to rank each gene in DE gene list. Gene sets relevant to pathways, processes and cell types of interest (blood-brain barrier, VEGF signaling, canonical Wnt/β-catenin signaling, non-canonical Wnt signaling, TGF-β signaling, extracellular matrix, cell-cell adhesion, transporters, Notch signaling, angiogenesis, endothelial cell proliferation, endothelial cell migration, apoptosis, antigen processing and presentation, inflammation, endothelial-to-mesenchymal transition, tip cells) were compiled as part of the overall gene set enrichment analysis, using the GO, KEGG, DAVID and GSEA databases as published previously.^68–70^ The ranked list of DE genes, along with the compiled gene sets, were loaded onto GSEA 4.1.0 software, and run through ‘GseaPreranked’ with the following settings: Number of permutations = 1,000; Collapse/Remap to gene symbols = No_collapse; Max size: exclude larger sets = 1,500; all other settings were left at ‘default’. Gene sets with FDR q value ≤ 0.3 were deemed significantly enriched and are included in the GSEA graphs.

### Statistical Analysis

Data presented as bar graphs indicate mean ± standard deviation (SD). Dots in bar graph represent individual values per mouse. Pairwise comparisons were calculated with unpaired or paired two-tailed t tests. Comparisons of one or more groups were performed using a 2-way ANOVA and p-values were adjusted based on the Holm Sidak post hoc correction. Statistical analyses were performed in GraphPad Prism (GraphPad Software, San Diego, CA, USA). Statistical details of experiments can be found in figure legends. Sample size was not predetermined using power analysis. Statistical differences between groups in Figure 8 are determined by Wilcoxon rank sum test comparing each group to the untreated control. Significance levels are indicated as follows: *p<0.05; **p<0.01; ***p<0.001; ****p<0.0001.

## DISCUSSION

FUS with microbubbles is a novel technique that allows non-invasive, localized, and transient increase in BBB permeability and presents a promising therapeutic method to deliver drugs for treatment of neurological and psychiatric diseases. However, the cell biological and mechanisms by which ECs respond acutely and sub-acutely to FUS are not fully characterized. By examining TJ morphology with confocal microscopy following FUS with microbubbles in two pressures, a low safe pressure associated with stable cavitation (450 kPa) used to deliver neurotrophic factors, and a high pressure associated with inertial cavitation (750 kPa) known to cause irreversible CNS damage,^30,40,71^ we find that BBB TJ integrity is compromised for a longer period (72 hours) at high, but not low, pressures in capillaries and to a lesser extent arterioles. The diameter of cavitating microbubbles matches more closely to that of capillaries than arterioles, allowing a higher probability of physical interactions with the vessel walls in smaller caliber vessels.^72^ This may explain why the microbubbles are more efficient to damage capillaries versus arterioles TJs at high pressure. Small and large vessels were equally likely to be permeable at low pressure whereas permeability of large vessels was favored at high pressures. With increasing vessel diameter, there is increased microbubble expansion^73,74^ and an increased inertial cavitation threshold,^75^ likely due to changes in the resonance frequency of the bubbles or additional constraints in the microvasculature.^76,77^ Thus, microbubbles may interact with the vessel wall in smaller vessels, inertial cavitation leading to microstreaming and shock waves events may occur more often in larger vessels,^7–11^ which is supported by our data that higher pressure damages more larger than smaller TJs. Although we could not detect any venules that colocalized with the fluorescent tracer at either pressure, structural changes in both transcellular (caveolae) and paracellular (TJs) BBB features have been observed in venules by immunoelectron microscopy at very high pressures (1 MPa), albeit at a lower percentage than capillaries and arterioles.^78^ Although BBB leakage may occur in venules at both low and high pressures tested in our study, the rate of diffusion may be very slow below the resolution limits of confocal microscopy. Future EM studies examining the low rate of transport in venules following FUS will provide better insight whether there is vessel selectivity in BBB opening with low safe pressures.

TJ segments are quite stable in the healthy CNS but undergo dynamic remodeling in ischemic stroke or Experimental Autoimmune Encephalomyelitis (EAE), an animal model for MS.^5,39,79,80^ Under conditions of stable cavitation, we find significant differences in both capillary and arteriole TJ segments with gaps larger than 2.5 μm between the ipsilateral and contralateral regions within 1 hour post FUS at both pressures that likely underlie FUS-mediated BBB opening. However, by 72 hours, there is no difference in the fraction of TJ segments with larger gaps (>2.5 μm) between the two hemispheres for 450 kPa, consistent with transient BBB opening. It has been reported that TJ protein degradation may occur as quickly as 20 minutes following sonication^38^ and TJ strands may be repaired within 1 hour post low pressure FUS. Our transcriptomic analysis of ECs within 1 hour post FUS revealed that “cellular responses to stress” and “junction protein synthesis” are upregulated to cope with injury; however, this response has resolved back to baseline levels by 72 hours post-FUS. This is consistent our findings that TJ abnormalities (i.e., TJ gaps >2.5 μm) are transient at low pressures in both capillaries and arterioles suggesting that TJ strands may undergo a transient reorganization within the membrane. Similar abnormalities were also seen with ZO-1 in Glut1+ vessels exposed to 450 and 750 kPa 72 hours after FUS (**Figure S12**). Stable cavitation may either change the turnover rate of TJ membrane protein or the phase transition state of ZO proteins that anchor TJ transmembrane proteins to the cytoskeleton.^81^ Under high pressure (750 kPa) with inertial cavitation, capillaries exhibited a full absence of TJ strands, a phenomenon likely induced by shock waves, generated by the collapse of microbubbles. These TJ deficits are not restored by 72 hours following sonication suggesting a permanent TJ damage. Transcriptomic analysis of ECs exposed to 750 kPa confirm cell biological findings that sustained injury (e.g. “cell death”) and repair processes (e.g. “cell proliferation”, “cell migration”, “vascular development”) were observed at both acute and long-term timepoints at 750 kPa. Removal of complete TJ segments may not permit ECs to restore cell-cell junctions by 72 hours leading to longer-term structural and functional BBB impairment at high pressure.

Could the transient opening of the BBB at low pressure mediated by caveolae? We observed no differences in BBB opening between *Tg::eGFP-Claudin5*^+/-^ mice and *Tg::eGFP-Claudin5*^+/-^; *Cav1^-/-^* mice sonicated at 450 kPa, suggesting that caveolae do not play a significant role in the transient BBB opening at safe pressures after 1 hour. Increased number of caveolae have been reported in arterioles using lower center frequency transducer after FUS^78,82,83^ and caveolae-mediated endocytosis may be important for transporting large molecules across the BBB.^84^ Although we cannot exclude the possibility that caveolae may play a role in transient BBB opening within the first hour following FUS at safe pressures, our data do not support a role for caveolar-mediated transport. The transcriptomic response of BBB opening in *Cav1*-/-mice should be investigated in future studies.

Although transient reactive microglial activation has been described after FUS,^85–87^ we find that microglia reside near the site of leakage along capillaries and arterioles one hour following FUS with microbubbles. However, by 72 hours microglia are indistinguishable from microglia on the contralateral hemisphere at low pressure, whereas they are present at a greater number and are highly phagocytic at high pressure. Microglia could respond to numerous factors in tandem following FUS, including the release of cytokines and chemokines,^20,88^ the secondary acoustic radiation force of oscillating microbubbles^89^ and serum components that have crossed the BBB.^90^ Our findings indicate the presence of fibrinogen, a known microglia activator, in the brain parenchyma after FUS. Fibrinogen deposition in the brain parenchyma decreases spine density and promoted cognitive decline.^91^ With pressures comparable to low FUS pressure used in the current study, no difference in neuronal excitability or dendritic tree morphology was observed between treated and untreated neurons at one week or 3 months following six weekly FUS sonications^92^ Moreover, FUS sonications targeting the striatum elicited no behavioral changes in the rotarod or open field test between treated and untreated groups for up to six months ^93^. Fibrinogen is pathogenic in multiple sclerosis,^53^ Alzheimer’s disease,^91,94^ brain trauma,^95^ and nerve injury.^96^ Fibrinogen was present in the brain parenchyma at both pressures one hour after FUS sonication. At 72 hours following low pressure FUS fibrinogen depositions were not present consistent with transient BBB opening. Interestingly, at 750 kPa 72 hours following FUS there is still BBB leakage and TJ proteins have not been fully restored in capillaries or arterioles; yet fibrinogen is not present in the brain parenchyma. The molecular weight of Fibrinogen (∼340 kDa) ^97^ is much larger than biocytin-TMR (840 Da), suggesting that BBB may have undergone some repair after 72 hours at higher pressure (FUS).

Similar to prior studies, we find no difference in GFAP intensity at either pressure at 1 hour, whereas there is increased GFAP intensity at 72 hours at both pressures. The activating factor(s) responsible for increased GFAP immunoreactivity are not clear. Numerous factors are known to cause astrocytes to adopt a reactive state including cytokines and growth factors, molecules released by cell injury or oxidative stress, as well as hypoxia and glucose deprivation^98^. After FUS, higher BBB permeability enhancement is associated with higher levels of inflammatory markers such as Cxcl1, Ccl2, Il1b, Il6, Tnf, and Icam1,^99^ which have been shown to alter the astrocyte phenotype into a more reactive state.^100^ At lower pressures many of these genes return to baseline by 24 hours,^101^ although reactive astrocytes may persist, even after injury resolution.^98^ It has been shown that GFAP immunoreactivity was elevated at 4 days, but returned to baseline by 15 days at safe FUS pressures.^85^ In contrast, at high FUS exposure levels GFAP immunoreactivity was present 7 weeks after a single sonication, and with morphological changes indicative of glial scar formation.^102^ Activated astrocytes can form a physical barrier to enclose an injured site and prohibit the passage of cytotoxic molecules into the brain parenchyma.^103,104^ At 750 kPa, astrocytes may act as a barrier to cytotoxic plasma components that could enter the brain through disrupted TJ strands. Reactive astrocytes can also inhibit axon regeneration,^105^ release neurotoxic factors^106^ and enhance synaptic degeneration.^107,108^ FUS at safe pressures has been shown to increase neurogenesis^109–112^ and FUS alone had no detrimental effect on synaptic density.^44^ Further studies investigating causal activators of astrocyte reactivity at different acoustic pressures are required to fully elucidate the mechanism of acute and long-term astrocyte reactivity after FUS and elucidate their contribution to BBB dynamic opening.

Evaluation of glial activation at later timepoints in response to various FUS-BBBO exposure conditions has also been reported by our group and others. Batts et al. characterized murine microglial and astrocyte activation following various FUS exposure conditions.^113^ This study reported a transient increase in Iba1 expression 24 hours post-FUS, and a delayed increase in GFAP expression 96 hours post-FUS across high and low FUS pressures. Moreover, Pouliopoulos et al., reported an increase in glial cell activation 2 days after FUS-BBBO in a rhesus macaque, however this activation was resolved and no gliosis was visible at 18 days following safe FUS exposure.^114^ Based on these studies we do not expect sustained glial activation in response to the pressures tested in this study, beyond the timepoints that were analyzed.

The activation of other support cells of the BBB by FUS-BBBO has also been reported and bears significance in interpreting the BBB functional deficits. Han et al., targeted the caudate putamen with high and low-pressure conditions resembling the high end of the exposures used in this study.^115^ Their low-pressure condition, which corresponds to a mechanical index of 0.62 MPa/MHz1/2 (compared to 0.37 and 0.61 MPa/MHz1/2 for our low and high exposure conditions, respectively) showed an increase in astrocytic Aqp4+ expression around vessels in the region of BBB disruption in both pressure conditions 48 hours after treatment. Thus, astrocytes respond to FUS-BBBO with structural modifications in their endfeet to encircle the barrier at the site of BBB disruption. This group also reported that GFAP expression was enhanced 48 hours after FUS exposure, suggesting that astrocytic swelling around the disruption site is a response mechanism to FUS-BBBO.

NVU transcriptomic changes that occur after FUS seem to facilitate viral transduction of microglia, astrocytes, endothelial cells, pericytes, oligodendrocytes and neurons 48 hours after exposure.^116^ Using lower MI values of 0.2 and 0.4, gene signatures indicative of resolving inflammation are observed in perivascular cells, such as pericytes. Therefore, NVU cells (pericytes and astrocytes), that are critical for the BBB, respond dynamically, and in a pressure-dependent manner to FUS-BBBO exposure. We anticipate that these cell types will respond similarly to the exposure conditions we report in the present study, assisting in BBB tight junction repair and functional reinstatement at the safe, lower pressure (450 kPa). Further studies investigating causal activators of astrocyte reactivity at different acoustic pressures are required to fully elucidate the mechanism of acute and long-term astrocyte reactivity after FUS and elucidate their contribution to BBB dynamic opening.

The transcriptomic analysis from single-cell RNAseq of brain ECs after FUS at both pressures revealed that all three EC subtypes (arterioles, capillary and venules) respond dynamically at the acute phase (1 hour) post FUS to upregulate transcripts related to injury processes (e.g. “cell death”, “inflammation”) and repair processes (e.g. “cell proliferation”, “cell migration”, “vascular development”) regardless of the applied pressure. Thus, ECs respond immediately to FUS injury to start repair processes. In contrast, there are distinct transcriptomic EC responses between the two pressures by 72 hours. Capillary ECs upregulate pathways related to “cell death”, “intracellular signal transduction,” “regulation of cell proliferation”, “vascular development” and “cellular response to stress” by 72 hours at both pressures. Surprisingly, GSEA analysis with targeted showed downregulated pathways for “blood-brain barrier”, “cell proliferation”, “angiogenesis”, “EC migration”, “tip cells”, “Wnt/β-catenin, Notch and TGFβ signaling” in ECs from 450 kPa by 72 hours. In contrast to 450 kPa, GSEA analysis revealed that processes such as “blood-brain barrier”, “inflammation”, “tip cells” were upregulated at 72 hour post-750 kPa, whereas those to “angiogenesis”, “EC migration”, VEGF, TGFβ and non-canonical Wnt signaling” were downregulated compared to controls. Moreover, transcriptome comparison of ECs from 750 kPa and 450 kPa revealed the presence of pathways related to “protein complex assembly,” “regulation of cell death,” and “response to wounding”. CNS ECs from 450 kPa are likely still undergoing vascular growth and BBB maturation at 72 hours post-FUS reflected in lower transcripts compared to controls. Moreover, our transcriptome findings are consistent with the concept that ECs exposed to 450 kPa mostly recover by 72 hours post-FUS, whereas those exposed to 750 kPa are still undergoing cell death and early repair processes, since they are damaged more extensively by high pressure.

The pathway analysis of EC transcriptome after FUS revealed that two angiogenesis promoting pathways TGF-β and Notch signaling change dynamically after FUS. TGF-β signaling which functions to inhibit cell proliferation, suppress inflammation and promote vascular development, is upregulated in capillary ECs one hour post-FUS together with inflammatory responses; However, it is largely downregulated by 72 hours relative to either control ECs, consistent with reduced inflammation and promotion of angiogenesis. Notch signaling is also upregulated significantly in capillary ECs one hour post-FUS; however, it is shut down by 72 hours post-FUS in capillary ECs, in particular for 750 kPa. Therefore, several signaling pathways critical for angiogenesis and EC proliferation change dynamically after FUS, but they remain lower in high pressure by 72 hours consistent with more blood vessel damage. Surprisingly, Wnt/β-catenin was not changed at the acute phase for both pressures, compared to the control group. However, canonical Wnt signaling was higher at the acute than the sub-acute (72 hours) phase post-FUS for both pressures and at lower than higher pressures by 72 hours consistent with continuous “EC proliferation” and “angiogenesis” at this time point at 750 kPa. Overall, this analysis demonstrates a reduction in the injury response and “return to normalization” at both pressures by 72 hours since Wnt/β-catenin signaling is reduced once angiogenesis and BBB maturation are complete.^1,2^

Finally, validation of the above single-cell RNA sequencing analysis was performed using a FISH assay labeling Egfl7 and Angpt2. The results of these assays are consistent with the sequencing data which demonstrate a significant upregulation of these markers 72 hours after FUS-BBBO at both high and low pressures. Comparison of Glut1+ vessels reveals greater expression of both Egfl7 and Angpt2 in the treated (ipsilateral) compared to control (contralateral) hemispheres of the brain in response to both 450 and 750 kPa.

Clinical trials currently employ lower frequencies compared to 1.5 MHz used in the current study. Using a lower frequency reduces aberrations caused by the skull and increases the focal volume. While well suited for human studies, lower frequencies pose challenges in small animal studies including the formation of standing waves creating peaks of high pressure associated with damage and heating.^117–119^ Microbubble diameter and the resonance frequency of the transducer will affect how microbubbles oscillate within the vasculature.^19,74^ As clinical trials utilizing FUS-mediated BBB opening for tumor therapy are under way,^23,120^ a rigorous understanding of NVU recovery will further aid in enhancing harnessing this technique for therapeutic treatment.

### Statistical Analysis

Data presented as bar graphs indicate mean ± standard deviation (SD). Dots in bar graph represent individual values per mouse. Pairwise comparisons were calculated with unpaired or paired two-tailed t tests. Comparisons of one or more groups were performed using a 2-way ANOVA and p-values were adjusted based on the Holm Sidak post hoc correction. Statistical analyses were performed in GraphPad Prism (GraphPad Software, San Diego, CA, USA). Statistical details of experiments can be found in figure legends. Sample size was not predetermined using power analysis. Statistical differences between groups in Figure 8 are determined by Wilcoxon rank sum test comparing each group to the untreated control. Significance levels are indicated as follows: *p<0.05; **p<0.01; ***p<0.001; ****p<0.0001.

## Acknowledgements

We thank Shutao Wang and Camilo Acosta for their supporting role in this work, as well as Tyler Cutforth for critical comments on the manuscript. T.K., R.L.N., R.J. and E.E.K. are supported by the National Institute on Aging of the National Institutes of Health under Award Number R01AG038961. D.A. is supported by the NIH (R01EY033994; R61/33 HL159949; RF1 AG078352, R21NS130265) and by an unrestricted gift to the Division of Cerebrovascular Diseases and Stroke in the Department of Neurology, CUIMC. The content is solely the responsibility of the authors and does not necessarily represent the official views of the National Institutes of Health.

