## Supplementary Information for "Safe focused ultrasound-mediated blood-brain barrier opening is driven primarily by transient reorganization of tight junctions"

**This PDF file includes:**

Figures S1 to S12

Legends for Figures S1 to S12

Legends for Single-Cell RNA-sequencing Tables S1-8

**Other supporting materials for this manuscript include the following:**

Single-cell RNA-sequencing Tables S1-8

**SUPPLEMENTARY INFORMATION**

**SUPPLEMENTARY FIGURES AND FIGURE LEGENDS**

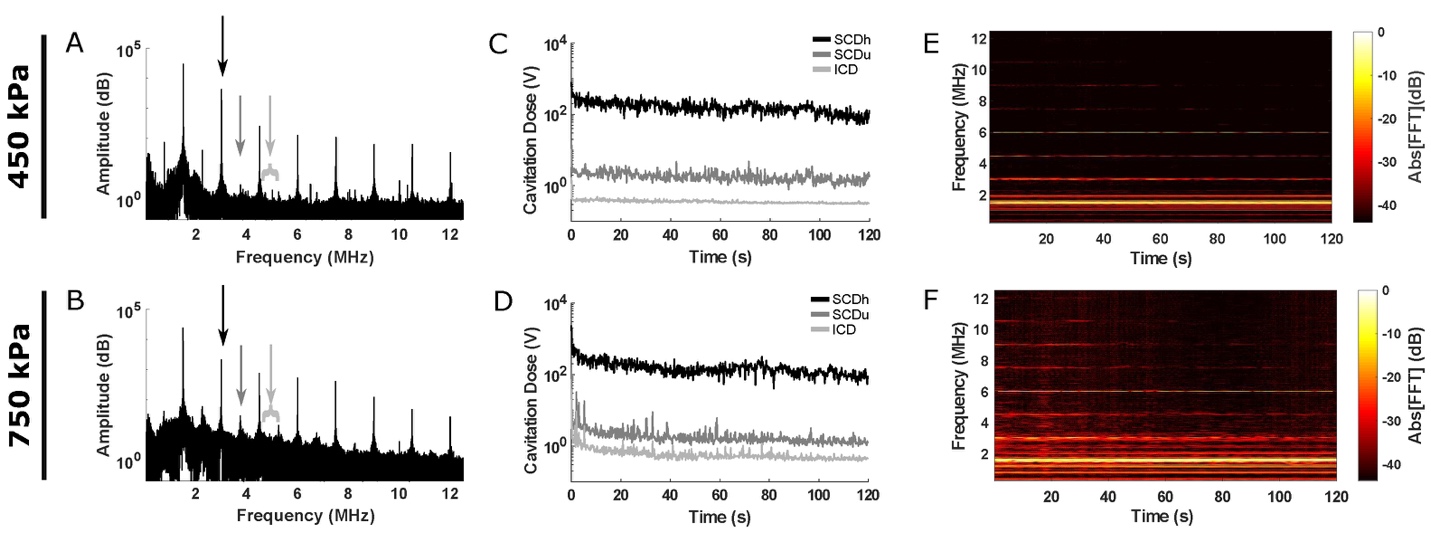

**Figure S1. Increased ultra-harmonic and inertial cavitation at 750 kPa confirmed by *in vivo* monitoring of passive cavitation.**

(**A, B**) The spectral amplitude for a single FUS pulse at 450 kPa (**A**) and 750 kPa (**B**) with black arrows indicating harmonic peaks (black arrow; nfc, n=1, 2… 6, fc=1.5 MHz), dark gray arrows indicating ultra-harmonic peaks (dark gray arrow; nfc/2, n=3, 5, 7, 9), and inertial cavitation defined as the broadband signal between harmonic and ultra-harmonic peaks (light gray). An increased amplitude of the ultra-harmonic peaks and broadband signal is seen at 750 kPa (**B**). (**C, D**) Sample traces of stable harmonic cavitation levels (SCDh, black), stable ultra-harmonic levels (SCDu, dark gray) and inertial cavitation levels (ICD, light gray) throughout the FUS sonication period (120 seconds) for 450 kPa (**C**) and 750 kPa (**D**). At 450 kPa, SCDu and ICD levels remain constant over the duration of the sonication, and there is a slight decrease in SCDh cavitation dose over time (**C**). At 750 kPa, both SCDu and ICD are highest at the beginning of the FUS sonication period until slowly reaching a plateau, suggesting some microbubble collapse (**D**). The inertial cavitation dose remains higher throughout the sonication at 750 kPa compared to 450 kPa. (**E, F**) Spectrograms for the sonication period (120 s) at 450 kPa (**E**) and 750 kPa (**F**). Broadband emissions indicated by signal intensity between harmonic and ultra-harmonic values are absent at 450 kPa (**E**), but present throughout the sonication period at 750 kPa (**F**).

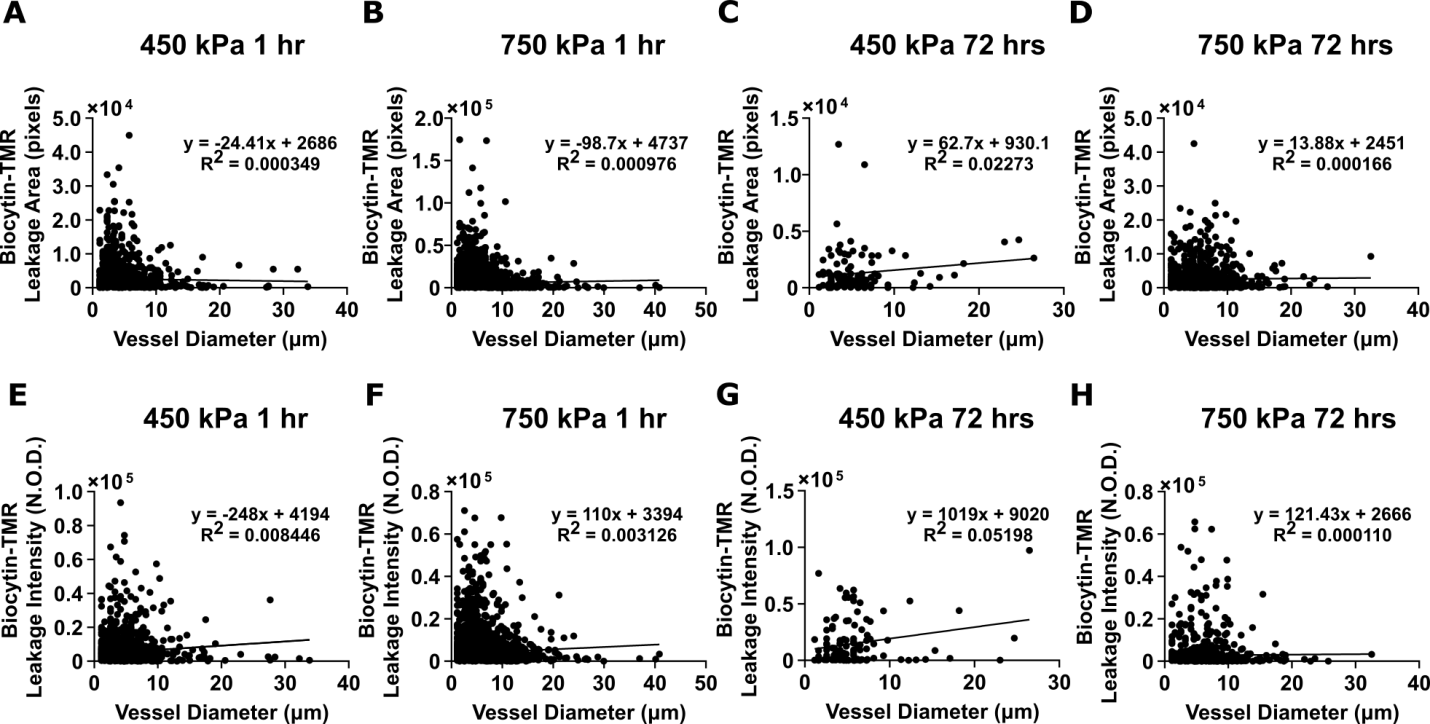

**Figure S2. The vessel diameter is a poor predictor of the area and the intensity of tracer leakage in either low or high pressures at 1 and 72 hours post-FUS with 1.5 MHz frequency.**

(**A-D**) Plots of the vessel diameter (X axis; μm) versus the area of biocytin-TMR leakage (Y axis) with the linear regression equation (formula and line) after 1 hour at 450 kPa (**A**; R^2^ = 0.000349) and 750 kPa (**B**; R^2^ = 0.000976; *p<0.05, one way ANOVA) and 72 hours at 450 kPa (**C**; R^2^ = 0.02273) and 750 kPa (**D**; R^2^ =0.000166). (**E-H**) Plots of the vessel diameter (X axis; μm) versus the intensity of biocytin-TMR tracer leakage (Y axis) with linear regression (formula and line) after 1 hour at 450 kPa (**E**; R^2^ = 0.008446, ***p<0.001, ANOVA) and 750 kPa (**F**; ****p<0.0001, R^2^ = 0.003126) or 72 hours at 450 kPa (**G**; *p<0.05, R^2^ = 0.05198) and 750 kPa (**H**; R^2^ =0.000110). For these dataset, the number of mice at 1 hour is N = 4 (n = 1387 vessels for 450 kPa and n = 4981 vessels for 750 kPa) and at 72 hours is N = 4 (n = 118 vessels for 450 kPa) and N = 5 (n = 940 vessels at 750 kPa). The slope of the linear regression was significantly different from zero for the vessel diameter versus the area of biocytin-TMR leakage at 1 hour for 750 kPa (**C**), the vessel diameter versus the intensity of biocytin-TMR leakage at 1 hour for 450 kPa and 750 kPa (**E, F**), and at 72 hours for 450 kPa. (**G**). Overall, the low R^2^ values of linera regression for all conditions suggest that vessel diameter is a poor predictor of vessel leakage for both pressures and time points analyzed in this study.

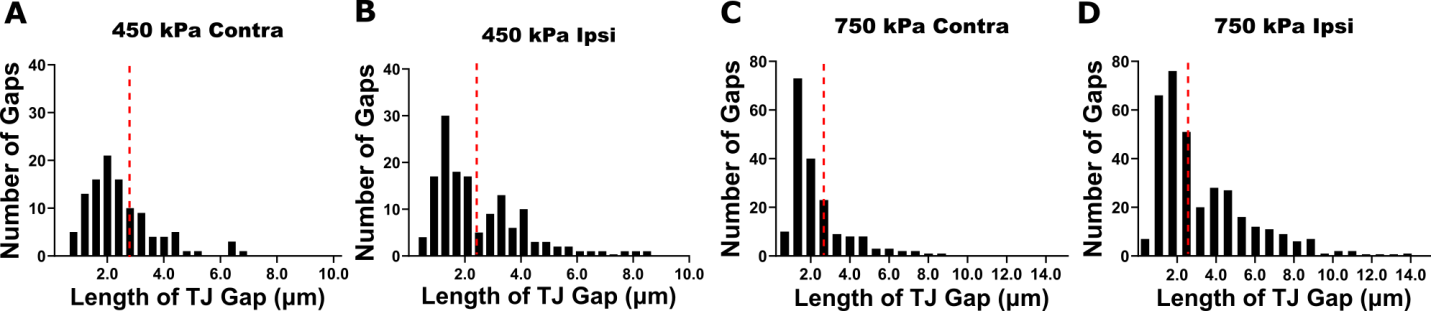

**Figure S3. Quantification of blood-brain barrier tight junction gap length one hour after FUS with 1.5 MHz frequency across blood vessels reveals a significant difference at the high, unsafe, pressure.**

(**A-D**) Histograms of the number of the blood-brain barrier tight junction strands with gaps (Y axis) plotted over tight junction gap lengths (X axis; µm) across CNS vessel types for all mice for 450 kPa (**A, B**) and 750 kPa (**C, D**) in the contralateral (**A, C**) and ipsilateral (**B, D**) hemispheres. On the contralateral hemisphere, 75% of the data are clustered below 2.9 μm at 450 kPa (red line) and 2.2 μm at 750 kPa (red line). Tight junction gaps > 2.5 μm in length were chosen as the cutoff for abnormal gap length in both datasets. Both 450 kPa and 750 kPa show a clear bimodal distribution in the length of tight junction gaps. The number of mice is N = 4 (450 kPa) and N = 5 (750 kPa) at 1 hour and the number of vessels is n = 254 (450 kPa) and n = 526 (750 kPa).

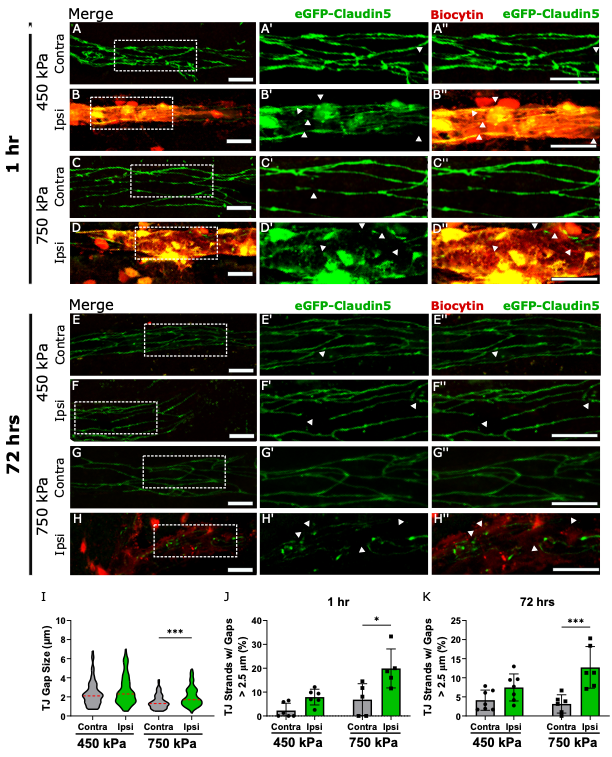

**Figure S4. Persistent structural abnormalities in arteriole tight junctions visualized by confocal microscopy correlate with continuous BBB opening at an unsafe pressure.**

(**A-D”; G-J”**) Maximum intensity projection images of 25µm-thick caudate putamen sections from *Tg::eGFP-Claudin5*^+/-^ mice 1 (**A-D”**) and 72 (**G-J”**) hours after 1.5 MHz FUS sonication at 450 kPa (**A-D”**)  and 750 kPa (**G-J”**) pressures in contralateral and ipsilateral hemispheres arterioles. eGFP (green) labels endothelial tight junctions and biocytin-TMR tracer leakage (red) is seen exclusively in the treated hemisphere within the brain parenchyma. TJ strand gaps are indicated by white arrowheads. (**E-F**) Quantification of the fraction of TJ strands containing gaps between 0.4-2.5 µm (**E**) and > 2.5 µm (**F**) at 450 kPa and 750 kPa one hour after FUS sonication. The fraction of TJ strands with gaps > 2.5 µm is significantly different between the ipsilateral and contralateral hemispheres at 750 kPa (N=4 mice; ^∗^p < 0.05; one-way ANOVA with Sidak’s multiple comparison test between contralateral and ipsilateral hemispheres). (**K-L**) Quantification of the fraction of TJ strands with gaps between 0.4-2.5 µm (**E**) or > 2.5 µm (**F**) at 450 kPa (N=4 mice) and 750 kPa (n=5 mice) 72 hours after FUS sonication. The fraction of TJ strands with gaps > 2.5 µm is significantly different between the ipsilateral and contralateral hemispheres at 750 kPa (N=4 mice; ^**^p < 0.01; one-way ANOVA with Sidak’s multiple comparison test between contralateral and ipsilateral hemispheres).  All data are presented as mean ± standard deviation. Scale bar=20 µm. See also **Figure S3 and S5**.

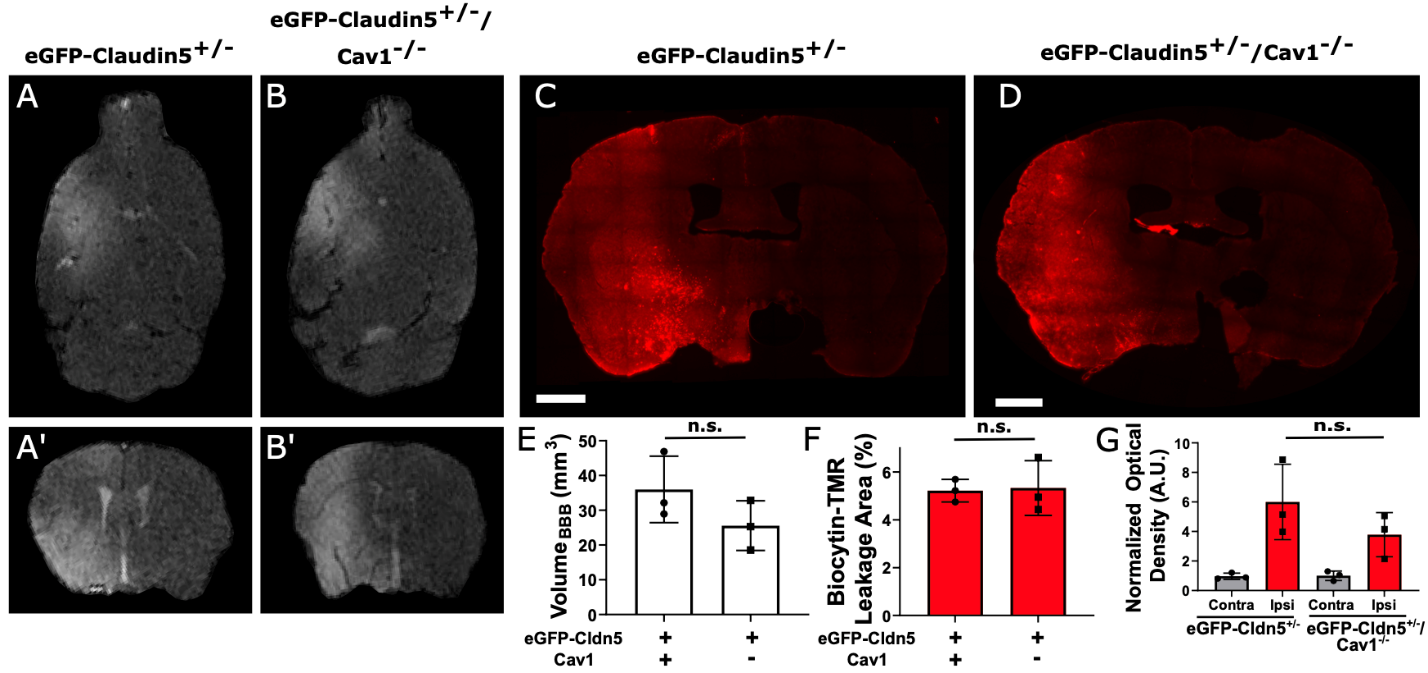

**Figure S5. Caveolin-1 is not required for the transient increase in BBB permeability following FUS with 1.5 MHz frequency at low safe pressures.**

(**A-B’**) Axial (**A, B**) and coronal (**A’, B’**) evaluation of increased BBB permeability 45 minutes-post FUS with microbubbles at 450 kPa with contrast-enhanced T1-weighted MRI in the caudate/putamen region of *Tg::eGFP-Claudin5+/-* mice (labelled as *eGFP-Claudin5+/-*) (**A, A’**) and *Tg::eGFP-Claudin5+/-; Cav1-/-* mice (labelled as *eGFP-Claudin5+/- / Cav1-/-*) (**B, B’**). (**C, D**) *eGFP-Claudin5+/-* and *eGFP-Claudin5+/-; Cav1-/-* mice were injected with biocytin-TMR (890 Da) tracer 1 hour after FUS with microbubbles to determine BBB permeability. Brain sections from *eGFP-Claudin5+/-* (**C**) and *eGFP-Claudin5+/-* / *Cav1-/-* (**D**) mice show a similar pattern of biocytin-TMR tracer leakage from blood vessels into the ipsilateral caudate/putamen region 1 hour following FUS-mediated BBB opening at 450 kPa. (**E**) Quantification of Gd-DPTA-BMA (540 Da) volume 45 min post-FUS at 450 kPa with matched VOIs between *eGFP-Claudin5+/-* and *eGFP-Claudin5+/-* / *Cav1-/-* mice (N = 3 mice/group). (**F**) Quantification of biocytin-TMR leakage area at 450 kPa one hour after FUS in *eGFP-Claudin5+/-* and *eGFP-Claudin5+/-* / *Cav1-/-* mice (N = 3 mice/group; n.s. = p>0.05 with Student’s t-test). (**G**) Quantification of biocytin-TMR fluorescence intensity normalized to optical density between ipsilateral and contralateral matched ROIs 1 hour after FUS at low safe pressures (450 kPa) (N = 3 mice/group; n.s. = p>0.05 with Student’s t-test between ipsilateral hemispheres; a.u., arbitrary unit). There is no difference in the area of intensity of tracer leakage between *eGFP-Claudin5+/-* and *eGFP-Claudin5+/-; Cav1-/-* mice one hour after FUS with microbubbles at 450 kPa suggesting a caveolar-independent mechanism for BBB opening at safe pressures. Scale bars are 1mm.

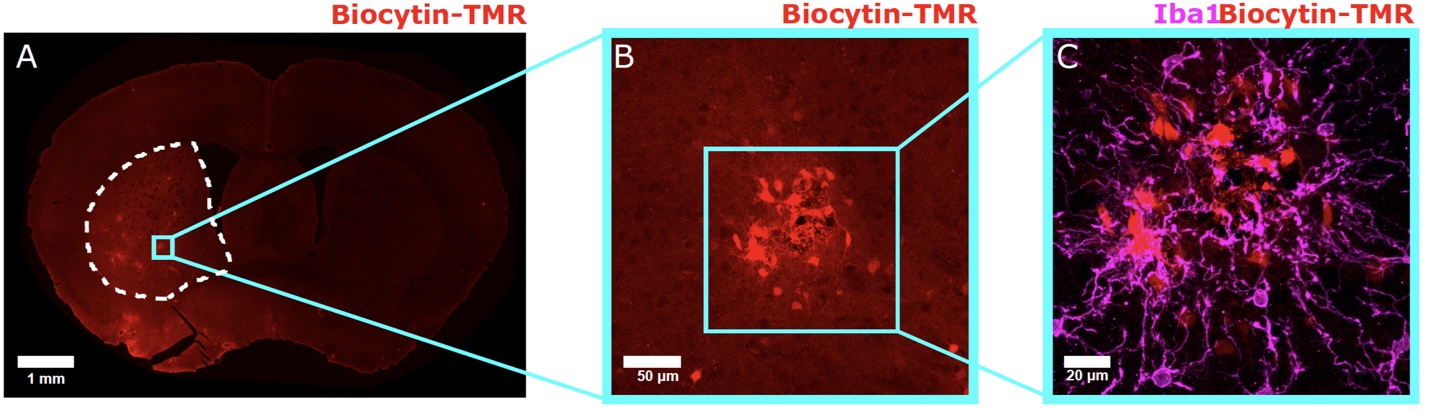

**Figure S6. Areas of biocytin-TMR leakage from blood vessels in the caudate/putamen that were chosen for microglia analysis.**

(**A**) Image of biocytin-TMR tracer leakage (red) from the brain vasculature into the ipsilateral hemisphere 1 hour after FUS-mediated BBB opening at 750 kPa. The areas of BBB leakage chosen for analysis were located within the caudate/putamen region outlined by the dotted white line. (**B, C**) Higher magnification images demonstrate regions of BBB leakage to biocytin-TMR (B; red) that were chosen for analysis of Iba1+ microglia /macrophages (**C**; Iba1 is in magenta).

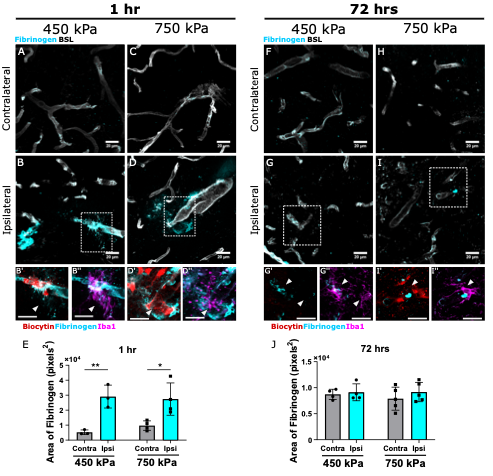

**Figure S7. Fibrinogen leaks across capillaries and arterioles only at the acute phase following FUS sonication with either safe or unsafe pressures.**

(**A-D”; F-I”**) Maximum intensity projection images of 25µm-thick caudate putamen sections from *Tg::eGFP-Claudin5*^+/-^ mice at 1 (**A-D”**) and 72 (**F-I”**) hours following 1.5 MHz FUS sonication at 450 kPa (**A-B”; F-G”**) and 750 kPa (**C-D”; H-I”**). Biocytin-TMR tracer (red) labels leaky vessels, lectin (white) labels vessels, Iba1 (magenta) labels microglia or macrophages and fibrinogen (cyan) labels the deposited serum protein into the CNS. (**E, J**) Quantification of the area of fibrinogen leakage at 1 (**E**) and 72 (**J**) hours following FUS sonication at 450 kPa (N=4 mice) and 750 kPa (N=5 mice) (*p<0.05 and **p<0.01; one-way ANOVA with Sidak’s multiple comparison test between contralateral and ipsilateral hemisphere). All data are presented as mean ± standard deviation. Scale bar=20 µm. **See also Figure S8.**

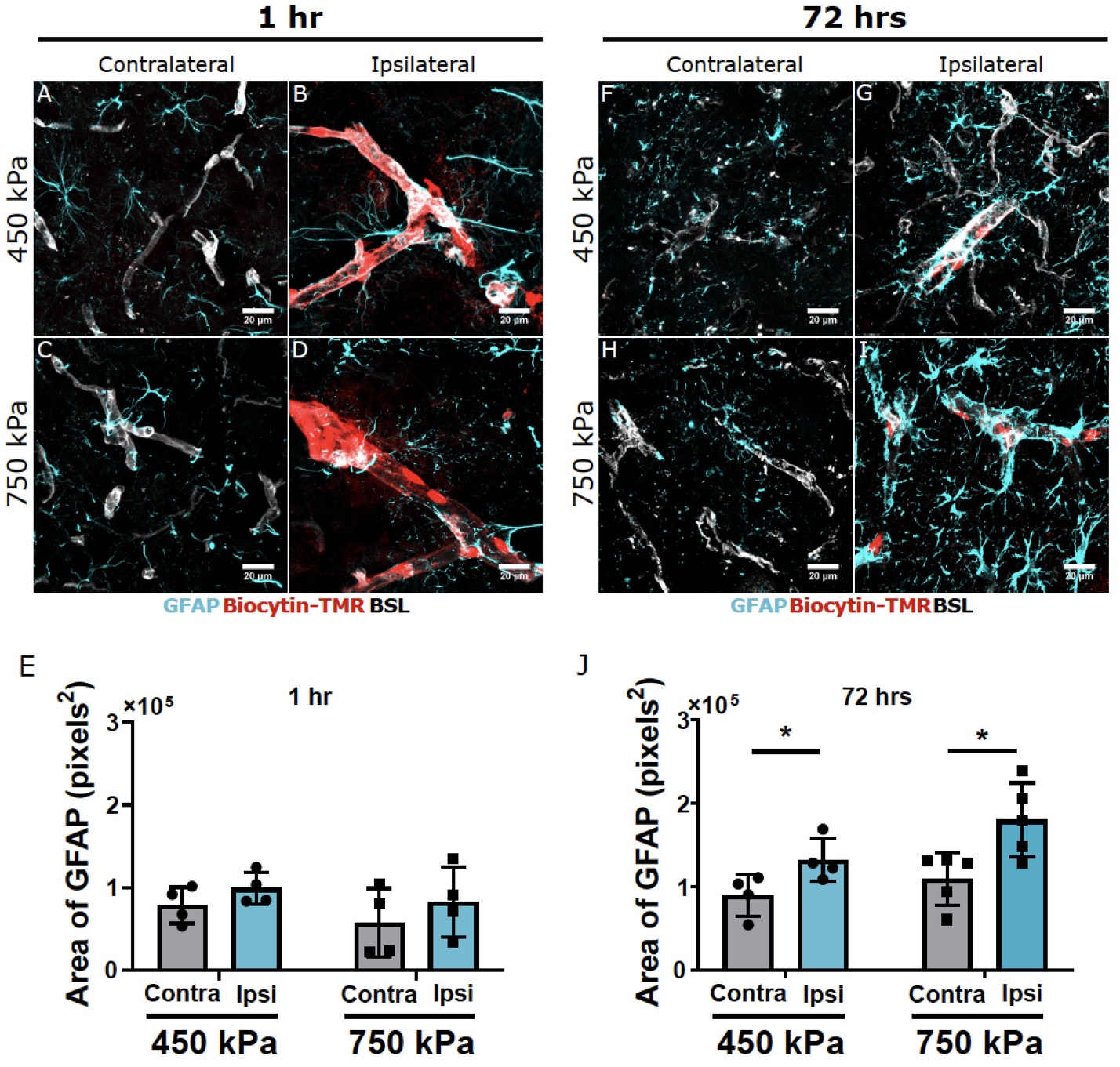

**Figure S8. Astrocyte reactivity is present in the caudate/putamen regions at 72 hours following FUS with 1.5 MHz frequency in both pressures.**

(**A-D; F-I**) Maximum intensity projection images of 25µm thick sections from the caudate/putamen region of *eGFP-Claudin5*^+/-^ brains at 1 (**A-D**) and 72 (**F-I**) hours following FUS sonication (1.5 MHz frequency) at 450 kPa (**A, B, F, G**) and 750 kPa (**C, D, G, I**). GFAP (cyan) labels normal and reactive astrocytes, biocytin-TMR tracer (red) labels blood vessel leakage, and BSL lectin (white) labels vessels. (**E, J**) Dotted bar graphs of GFAP^+^ area at 1 (**E**) and 72 (**J**) hours following FUS sonication at 450 kPa (N = 4 mice) and 750 kPa (N = 5 mice) (*p<0.05; one-way ANOVA with Sidak’s multiple comparison test between the contralateral and ipsilateral hemisphere). Dotted bar graphs show mean ± standard deviation. Scale bars = 20 µm.

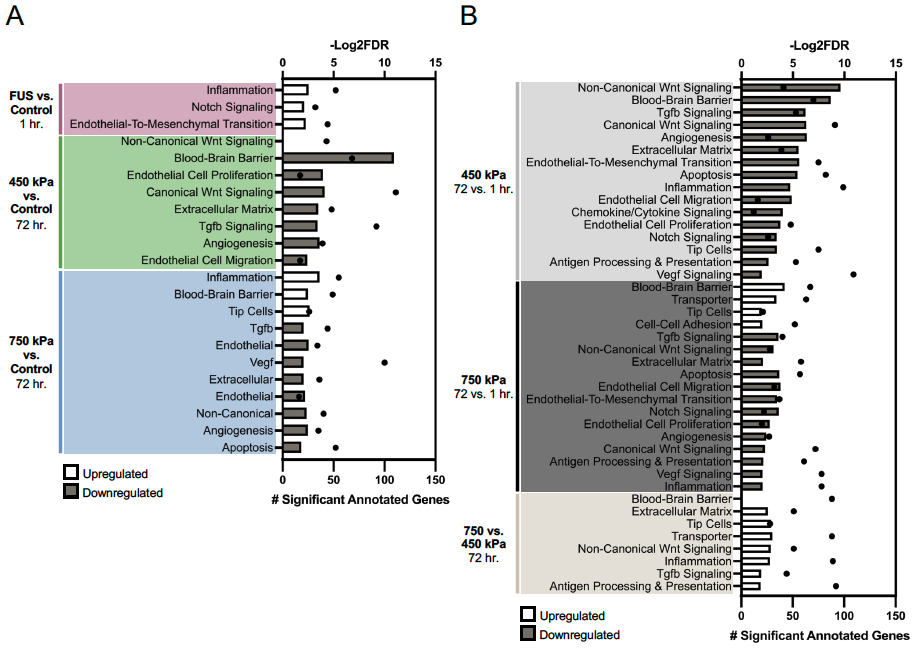

**Figure S9. Gene set enrichment (GSEA) results for capillary ECs.** (**A**) GSEA terms up and downregulated in each pressure and timepoint condition relative to control (P value cutoff < 0.3) in capillary ECs. (**B**) GSEA terms up and downregulated between 72 and 1 hour within each pressure group, then between 750 and 450 kPa at 72 hours post-FUS-BBBO (P value cutoff < 0.3). See also **Table S8**.

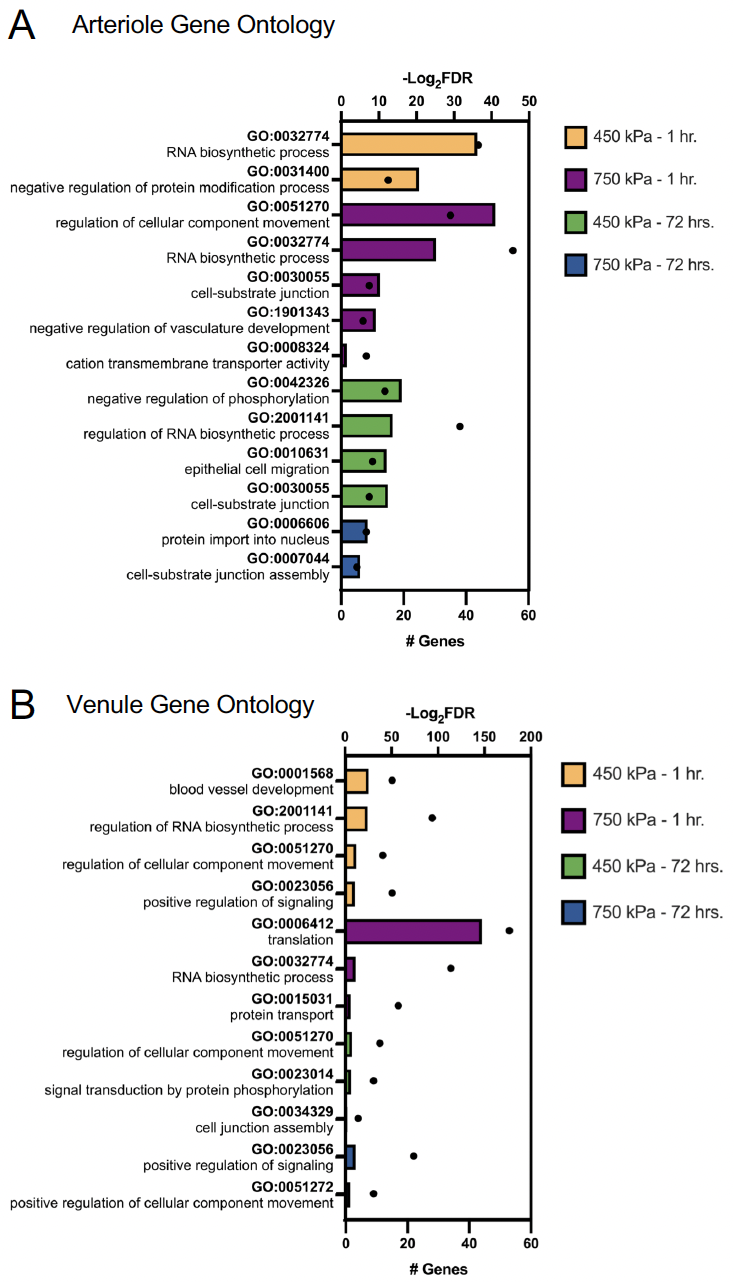

**Figure S10. Gene ontology analysis of differentially-expressed genes from the single cell RNA-sequencing data of brain arteriole and venule endothelial cells isolated after 1 and 72 hours post-FUS at both pressures.**

(**A, B**) Upregulated gene ontology terms of differentially expressed genes from the single cell RNA-sequencing data in arteriole (**A**) and venule (**B**) ECs for each of the four treatment groups relative to untreated controls. Bars represent -Log_2_(FDR) values (top axis), and dots represent the number of significantly differentially expressed genes annotated to each term (bottom axis).

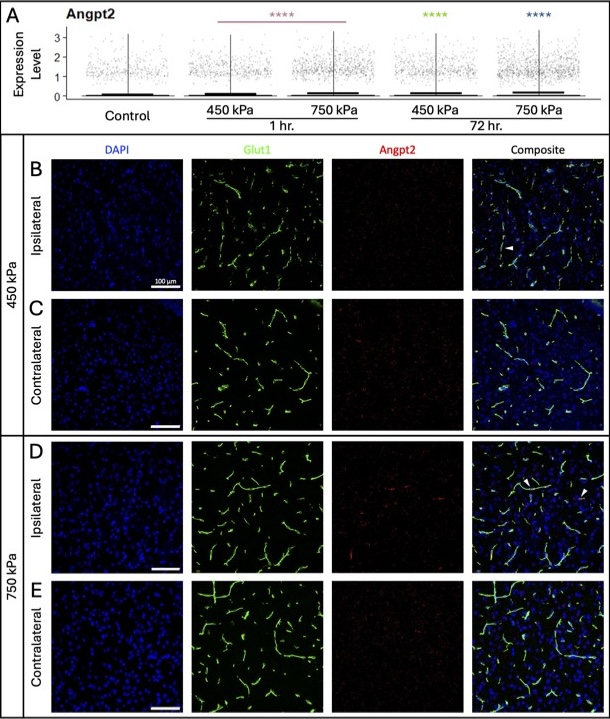

**Figure S11**. Glut1-positive endothelial cells exposed to 750 kPa exhibit a sustained, FUS-induced increase in Angpt2 expression by fluorescence in situ hybridization assay. (A) Violin plots showing expression of Angpt2 across each of the treatment groups. Mean expression level is indicated by the bar in each plot. Statistically significant differences between each group and untreated controls are determined by Wilcoxon rank sum test. **** P ≤ 0.0001. EC’s exposed to 450 kPa (B) exhibit heightened levels of Angpt2 expression compared to the untreated contralateral hemisphere (C). EC’s exposed to 750 kPa (D) exhibit markedly greater expression of Angpt2 relative to the untreated contralateral hemisphere (E). White arrowheads indicate Glut1-positive vessels with positive Angpt2 expression.

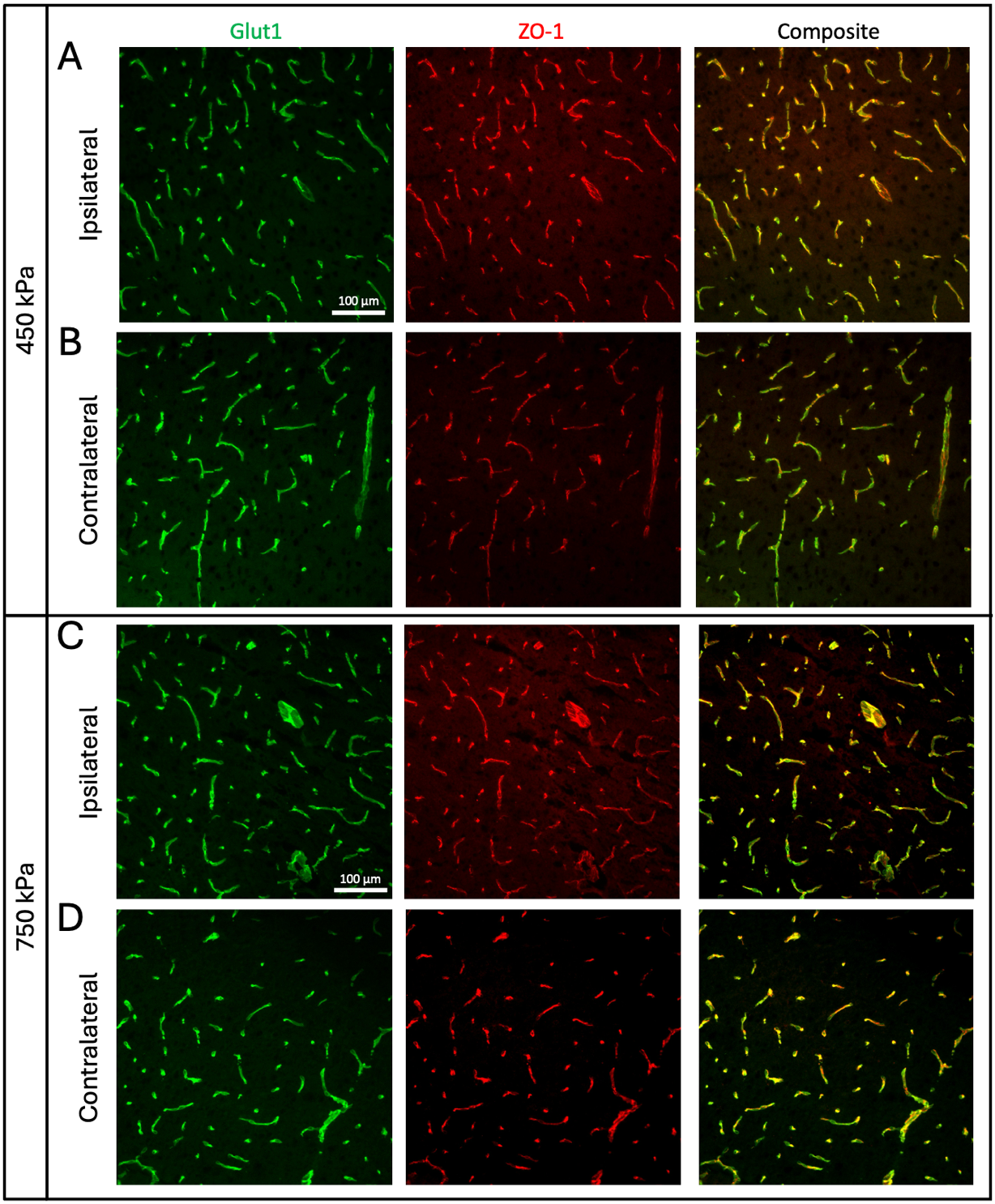

**Figure S12**. ZO-1 expression after high and low FUS-BBBO exposure. The colocalization of ZO-1 with Glut1+ vessels is shown on the ipsilateral (A) and (B) contralateral hemispheres of mice treated with 450 kPa FUS. The colocalization of ZO-1 with Glut1+ vessels is shown on the ipsilateral (C) and (D) contralateral hemispheres of mice treated with 750 kPa FUS.

**SUPPLEMENTARY DATASET**

**TABLES S1 – S8. Results of the single-cell RNA-sequencing analysis of brain ECs from the FUS experiments.**

Batch structure for the single-cell RNA-sequencing experiments (**S1**). Table of differential gene expression between control vs. FUS 1 hour at both pressures (**S2**); control vs. FUS 72 hours at 450 kPa (**S3**); control vs. FUS 72 hours at 750 kPa (**S4**). Negative log2fc values mean that the gene is lower in the FUS compared to control, and positive values mean that the gene is higher in FUS compared to control. Go analysis generated from the DEGs for the capillary (**S5**), arterial (**S6**) and vein (**S7**) ECs. GSEA analysis results are summarized for each condition (**S8**).
